# OT-Curtains: an approach for studying protein interactions with DNA-ends using optical tweezers and confocal fluorescence microscopy

**DOI:** 10.64898/2025.12.05.692558

**Authors:** Sara De Bragança, Clara Aicart-Ramos, Ángel Rivera-Calzada, Raquel Arribas-Bosacoma, Mark S. Dillingham, Oscar Llorca, Fernando Moreno-Herrero

## Abstract

DNA ends generated by double-strand breaks are vulnerable intermediates that must be rapidly recognized, protected and resolved to preserve genome integrity. We present OT-Curtains, a single-molecule method inspired by DNA Curtains that uses a custom branched DNA substrate containing multiple accessible ends for simultaneous observation on dual-trap optical tweezers coupled to confocal fluorescence microscopy. Eliminating DNA surface anchoring, facilitating rapid protein and buffer exchange, and offering force-free experiments, OT-Curtains overcomes common limitations of flow-stretch-based methods. OT-Curtains allows real-time visualization and quantification of end recognition, protection, resection and cleavage at several DNA ends in parallel. We demonstrate compatibility with well-studied DNA binding systems by monitoring Ku-mediated DNA break recognition, AddAB-mediated DNA break resection, ParB-mediated DNA condensation and KpnI-mediated DNA cleavage. We show that kinetic and mechanistic parameters can be extracted from the data under defined forces and solution conditions. OT-Curtains offers an accessible and multiplexed route to interrogate DNA-end transactions central to double-stranded DNA break repair pathways and telomere biology, as well as a general framework for benchmarking proteins acting at DNA ends.

## Introduction

In cells, genome integrity is safeguarded by the coordinated action of molecular machineries that organize, replicate, and repair chromosomal DNA. Because the DNA is continually engaged by these essential biological processes, it remains highly vulnerable to damage. Among the various types of DNA lesions, double-stranded DNA breaks (DSBs) are particularly cytotoxic because they can be lethal to the cell if left unrepaired or can drive genome instability if misrepaired. DSBs arise both physiologically, as is the case with programmed DSBs during meiotic recombination and V(D)J recombination, or pathologically as a consequence of replication stress, metabolic byproducts, or exposure to exogenous DNA damaging agents such as ionizing radiation and certain chemotherapeutic agents.

Exposed DNA ends are highly susceptible to degradation or aberrant ligation. To prevent this, the cellular DNA damage response deploys a diverse set of proteins that recognize, protect, and process DNA ends to ensure their accurate and controlled repair. For example, the DNA end-binding Ku70-Ku80 complex (named simply as Ku hereafter), rapidly associates with free DNA ends, stabilizing them, protecting them against nucleolytic degradation, and initiating repair through non-homologous end joining (NHEJ) ^1–3^. The MRN complex, in turn, acts together with resection factors to remodel DNA ends, facilitating repair through homologous recombination (HR) ^4–6^. In a distinct context, at telomeres, chromosome termini must be protected to prevent inappropriate recognition and processing. Specialized protein assemblies such as shelterin mask telomeric ends from DNA damage sensors ^7,8^. Understanding how proteins engage with and process DNA ends is crucial to elucidating the mechanisms behind genome stability and cellular aging, as well as to uncover how defects in these processes contribute to disease and oncogenesis.

Single-molecule techniques have revolutionized the study of protein-nucleic acid interactions by enabling direct visualization of molecular dynamics with high spatial and temporal resolution. Among these approaches, several have been specifically developed to investigate how protein factors engage with accessible DNA ends. For instance, using Magnetic Tweezers (MT), it has been possible to investigate how the helicase/nuclease AddAB unwinds and degrades DNA starting from an accessible end ^9–11^. Additionally, the fluorescence-based single-molecule Fluorescence Resonance Energy Transfer (smFRET) technique, employing DNA molecules labelled with Cy3 and Cy5, has allowed precise monitoring of DNA end joining by the NHEJ machinery ^12,13^, owing to the distance-dependence of the FRET signal.

A widely used class of single-molecule methods for studying protein interactions with DNA ends is based on the stretching of DNA molecules anchored to a surface or to an optically trap bead by laminar flow ^14–20^. This flow-stretching approach, combined with total internal reflection fluorescence microscopy (TIRFM) or epifluorescence, allows direct visualization of labelled DNA molecules and the proteins acting upon them. The introduction of microfabrication techniques and the use of a fluid lipid bilayer to tether the DNA molecules to a surface facilitated the alignment of hundreds of molecules, giving rise to the high-throughput DNA Curtains method ^17–19^. Despite their strengths, flow-stretched DNA methods face limitations that hinder broader application. Primary constraints are the requirement for specialized nanofabrication and the steep learning curve associated with mastering the DNA tethering strategy. In addition, TIRF-based methods encounter additional challenges regarding photodamage and the use of high protein concentrations, where increased background fluorescence can lower the signal-to-noise ratio and obscure protein-DNA interactions. Additionally, the need to tether DNA molecules to a surface, can introduce surface-related effects, such as nonspecific protein adsorption or undesired DNA binding. More generally, a common limitation of flow-stretched methods is that the DNA experiences a stretching force due to hydrodynamic drag that is dependent on its distance from the surface and minimal at the free DNA end^21^. This could induce artefacts in the apparent specificity of the system being studied in circumstances where the protein activity (binding and/or catalysis) differs depending on the degree of stretching^22,23^. Finally, rapid buffer exchange and exposure to different experimental conditions is challenging ^24^.

Here, we present a single-molecule approach inspired by DNA Curtains that offers similar advantages to the above-mentioned techniques but eliminates the surface-related limitations and simplifies protein and/or buffer exchange. We introduce Optical Tweezers (OT)-Curtains, an approach in which a custom-designed DNA molecule, when optically trapped at its termini, retains four short or long DNA branches extending from its backbone. We developed this method for a dual-trap optical tweezers combined with confocal fluorescence microscopy and microfluidics, providing a new use of a commercial system that allows the direct visualization of protein interactions with several DNA ends in parallel. The dedicated microfluidics system enables flow-stretching of the branches (if needed) and rapid exchange of reagents and experimental conditions. Our method allows direct visualization of multiple DNA ends simultaneously, facilitating the investigation of biological processes that occur at DNA ends, such as those involved in NHEJ, or DNA end resection by helicases and nucleases involved in HR. Moreover, we show that OT-Curtains can be applied to investigate other biological processes, including restriction activity without loss of the tethered molecule and DNA condensation by chromatin-associated proteins. OT-Curtains also represent a promising tool to investigate processes beyond the scope of this work, such as mismatch repair, the resolution of stalled replication forks, or interaction with single-stranded DNA binding proteins.

## Results and discussion

### The OT-Curtains Construct

To study biological processes involving accessible DNA ends using single-molecule techniques we developed OT-Curtains. This method integrated the concept of individually tethered DNA molecules with spatially organized, parallel DNA branches, each presenting a defined DNA end and sequence. The design is versatile and can be customized to address diverse biological questions.

At the core of the method is the OT-Curtains backbone plasmid, a 22,142 bp double-stranded DNA (dsDNA) containing four poly-BbvCI regions spaced approximately 3.7-4.2 kbp apart. Each region comprises five closely-spaced BbvCI restrictions sites separated by 15-16 bp. A detailed description of the OT-Curtains backbone plasmid construction is provided in the Methods section. To assemble the OT-Curtains construct, the plasmid was first linearized by BssHII digestion (**Fig. 1a**). Following plasmid linearization, site-specific nicking by Nt.BbvCI created five nicks at each poly-BbvCI region, all located on the same DNA strand (**Fig. 1b**). Subsequent heat treatment released the four 15-16 nts fragments from each region, producing 63 nt singlestranded gaps (**Fig. 1c**). These gaps can then be filled in with DNA structures containing sequences homologous to the gap region (**Fig. 1d-f**). Currently the OT-Curtains backbone substrate generates four identical gaps, but further modifications could be implemented to generate gaps with different sequences allowing the ligation of four fully independent branches, as illustrated in **Figure 1f**. To create two attachment points at the termini of the OT-Curtains backbone, a ∼1 kbp biotinylated DNA handle was produced by PCR through biotin-dUTP incorporation. Finally, the branch fragments (when required) and handles were ligated to the DNA backbone using T4 DNA ligase in an overnight, single tube reaction, yielding a ready-to-use biotin-tagged OT-Curtains construct.

**Fig. 1.**
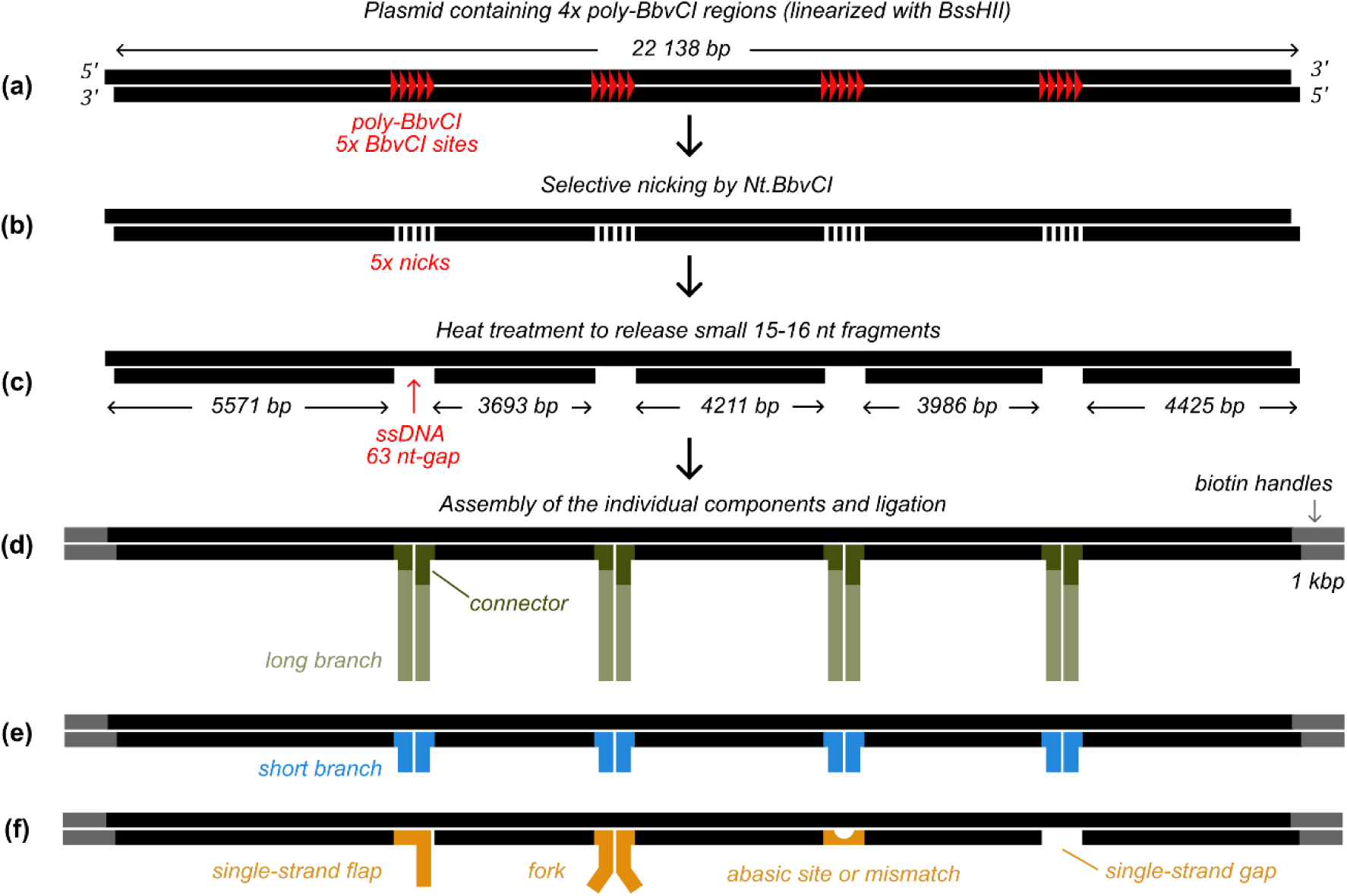
OT-Curtains substrate schematics. **(a)** Plasmid containing four poly-BbvCI regions. **(b)** Site-specific nicking by Nt.BbvCI creates five nicks per poly-BbvCI region. **(c)** OT-Curtains backbone molecule with four 63-nt single-stranded DNA gaps. **(d-f)** The OT-Curtains construct is assembled by incorporating biotinylated handles and branch structures of choice to the backbone. Variants addressed in this work have either long branches (**d**) or short branches (**e**), but different structural elements can be incorporated (**f**) depending on the biological question to be addressed.

To study DNA-end binding proteins, we built constructs containing either long-branches with specific sequences or short blunt-ended branches. In the former case, each branch consisted of two components: a connector and a long fragment. The connector, produced by annealing two synthetic oligonucleotides, was designed to hybridize to the 63-nt ssDNA gaps on the backbone. The long fragments were produced by PCR amplification using high-fidelity polymerases and the sequence templates of interest. Before annealing to the backbone, long fragments were joined to the connectors and gel purified to prevent competition from unbound connectors in solution (**Fig. 1d**). In the latter case, the short-blunt DNA structures were assembled by annealing synthetic oligonucleotides either to each other or directly to the 63-nt ssDNA gaps generated at the poly-BbvCI regions, thereby filling them (**Fig. 1e**). Substrate variants containing flaps, tails, abasic sites, or other modified structures can be created following this same strategy (**Fig. 1f**). In addition, the OT-Curtains substrate can also be adapted for applications requiring several short ssDNA regions (**Fig. 1f**). In this case, the substrate is ready to use immediately after handle ligation. If shorter gaps are required, synthetic oligonucleotides can be annealed within the 63nt gaps to reduce their length.

The modular design of the OT-Curtains construct enables an easy modification of both branch length and sequence, as well as the incorporation of fluorescent labels, specific overhangs, or complex DNA end structures to address specific questions related to the biological system under study. Notably, tandem molecules containing up to eight branches can also be obtained by adjusting the handle-to-backbone ratio (**Supp. Fig. S1**). This approach therefore introduces parallelization into the otherwise single-molecule optical tweezers technique. In this work, we employed short branches of 20 bp or 60 bp (sometimes carrying fluorophore modifications) and long branches of approximately 7 kb, containing a *parS* site when required. A detailed description of the fabrication and assembly of each type of substrate used in this work is provided in the Methods section.

### Proof-of-principle of substrate fabrication using end-labelled branches

To introduce the OT-Curtains method, we designed a protein-independent assay in which the branch termini were specifically labelled for detection by confocal fluorescence microscopy. The OT-Curtains construct with long branches consists of a DNA backbone with four ∼7 kbp branches, each labelled at its distal end with an ATTO488 fluorophore (**Fig. 2a** and **Suppl. Video 1**). Because the branches are relatively long, they must be extended to allow proper visualization. To achieve this, we applied a flow perpendicular to the main DNA axis, i.e., along the direction of the branches. Staining the DNA with 100 nM Sytox Orange (SxO) enabled simultaneous visualization of the DNA backbone and branch extensions, while also allowing detection of the fluorophore-labelled branch ends (**Fig. 2b**).

**Fig. 2.**
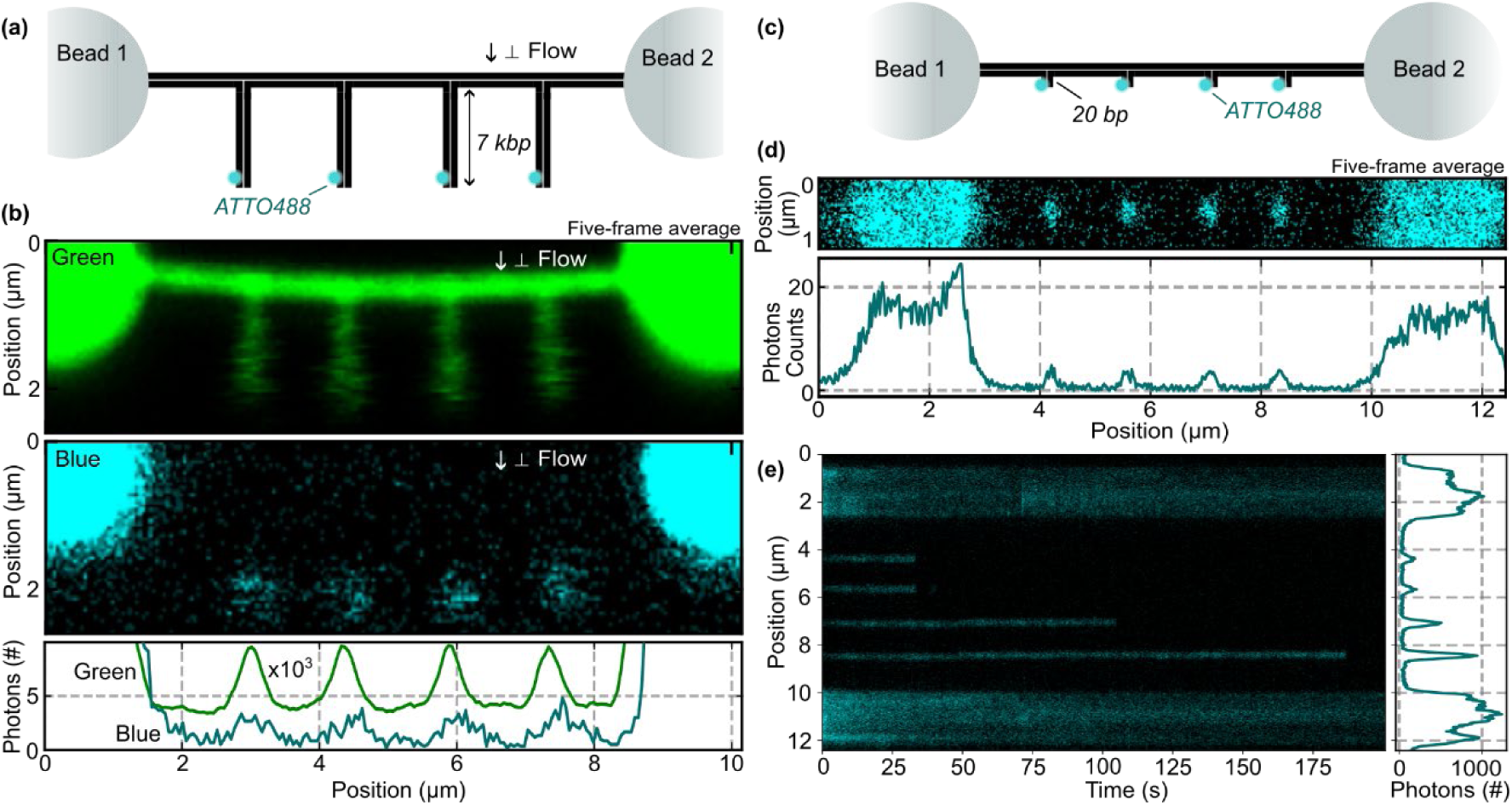
OT-Curtains schematics. **(a)** DNA construct with four long (∼7 kbp) branches, each labelled at its distal end with an ATTO488 fluorophore. **(b)** Two-dimensional confocal scan of the labelled construct shown in (a). The upper panel shows photon count data for Sytox Orange (SxO) emission acquired with the green filter. The middle panel shows photon count data for ATTO488 emission acquired with the blue filter. Images represent the average of five consecutive time frames. The lower panel shows photon count intensity summed along the y-axis. Note the three order of magnitude difference between the SxO (green) and ATTO488 (blue) photon counts. **(c)** DNA construct containing four short (20 bp) branches, each labelled at its distal end with ATTO488. **(d)** Two-dimensional confocal scan of the labelled construct shown in (c). Photon count data corresponds to the blue filter. Image represents the average of five consecutive time frames. **(e)** One-dimensional confocal scan (kymograph) along the DNA axis as a function of time, showing ATTO488 photobleaching. See **Supp. Fig. S2** for branch photon emission profiles.

In this configuration, experiments are typically performed under a constant flow to extend the branches. However, unlike near-surface single-molecule assays, where hydrodynamic drag is affected by wall effects, our configuration operates in bulk fluid, resulting in more predictable flow-induced forces in the order of 10^-2^ pN, with presumably little effect on protein activity. Indeed, a protein bound to the end of a branch in a flow generated by a pressure difference experiences a hydrodynamic drag force (*F*_*y,protein*_) that follows Stokes’ law and can be estimated from the local flow velocity (𝜈).

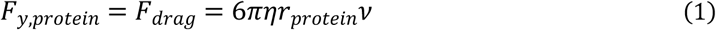

where *η* is the medium viscosity and *r*_*protein*_ is the protein hydrodynamic radius. The flow velocity can be calculated from the force measured on the trapped bead in the y-direction:

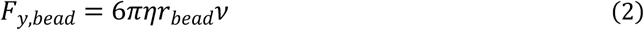

where *r_bead_* is the bead radius? Combining equations (1) and (2) yields a direct relation between the measured force in the trapped bead and the force experienced by the protein:

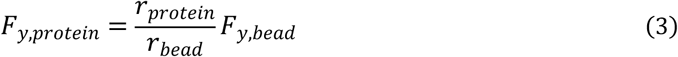

As an example, let’s consider a perpendicular flow that produces a 10 pN force in the y-direction on a 1.76 µm-diameter bead, and the AddAB heterodimer (studied below) bound to the end of an OT-Curtains long branch. AddAB is a globular protein complex with a molecular weight of ∼ 290 kDa when complexed with DNA ^25,26^. Using HullRad algorithm^27^, a translational hydrodynamic radius of 5.64 nm can be calculated from AddAB’s crystal structure. Under these conditions, the AddAB complex bound to the DNA end experiences a drag force of approximately 0.06 pN, about 100-fold smaller than the force acting on the bead, and has little to no effect on AddAB activity as assessed from MT experiments ^9,11^.

For force-free experiments that require a DNA end but not a long fragment, OT-Curtains with short branches is the optimal configuration (**Fig. 2c**). Because the branches are short, the fluorophore emission can be detected without applying a perpendicular flow to extend them, thereby ensuring true force-free conditions (**Fig. 2d**). Moreover, performing one-dimensional (1D) scanning along the DNA molecule improves temporal resolution and allows experiments to be maintained over extended periods (**Fig. 2e**, **Supp. Fig. S2**). In this configuration, staining the DNA with SxO is unnecessary, making it particularly suitable for applications where the use of DNA intercalating dyes should be avoided. Quantifying photon emission from each branch allows the estimation of the number of molecules bound at the DNA ends. Photon emission profiles from the branches shown in **Fig. 2e** indicate that a single ATTO488 fluorophore emits approximately 2-4 photons per scan line over an area of 10 pixels of 25 nm x 25 nm (**Supp. Fig. S2**).

In the following sections, we present results obtained using the OT-Curtains platform to investigate a range of biological systems. These results demonstrate the versatility of OT-Curtains for studying DNA-end binding processes such as DNA unwinding and resection occurring in homologous recombination, the binding of proteins involved in NHEJ, and other processes that might not require a DNA end such as DNA condensation and restriction.

### DNA-end resection by the helicase-nuclease AddAB

AddAB is a heterodimeric helicase-nuclease complex that initiates homologous recombination in bacteria by processing dsDNA ends ^28,29^. Upon binding a DNA end, AddAB couples ATP-driven unwinding to coordinated nuclease cleavage that is regulated by the presence of specific DNA sequences called Crossover hotspot instigator (Chi) sequences, generating a long 3′-terminated ssDNA tail suitable for RecA loading and strand exchange ^28^. The AddA subunit contains a 3′ to 5′ SF1 helicase motor domain and an associated nuclease domain, while AddB contributes a helicase-like module responsible for Chi recognition and a complementary nuclease domain. Together, the two subunits track along dsDNA, channelling the unwound products through separate paths ^30^. This architecture enables rapid, processive translocation coupled to tuneable resection activity, modulated by the presence of Chi sequences ^9,31^, cofactors (ATP, Mg^2+^), DNA-end structure, and protein roadblocks. Through tight coordination of unwinding and degradation, AddAB determines the length of resection, thereby setting up the substrate for downstream steps of HR. Beyond its role in DSB repair, AddAB also participates in replication fork restart by processing regressed forks and re-establishing replication templates ^28,32^.

To evaluate AddAB activity using OT-Curtains, we opted for a substrate variant containing four 7 kbp-long branches lacking Chi sequences and terminating in blunt ends (**Fig. 3a**; see Methods for details on fabrication). Upon exposure to 10.5 nM AddAB in the presence of 250 µM ATP, the lengths of the SxO-labelled branches progressively decreased (**Fig. 3b** and **Suppl. Video 2**). Because SxO specifically intercalates into double-stranded DNA, this shortening indicates progressive duplex unwinding and degradation, consistent with the combined helicase and nuclease activities of AddAB^33^. Within less than one minute, the AddAB complex unwinds the duplex while cleaving the emerging single-stranded DNA products. In the absence of ATP, AddAB bound stably to the DNA ends, as directly visualized using Quantum Dot-labelled AddAB (Qdot525-AddAB), and remained still at the DNA ends (**Fig. 3c** and **Suppl. Video 3**), demonstrating the strict ATP dependence of the unwinding process.

**Fig. 3.**
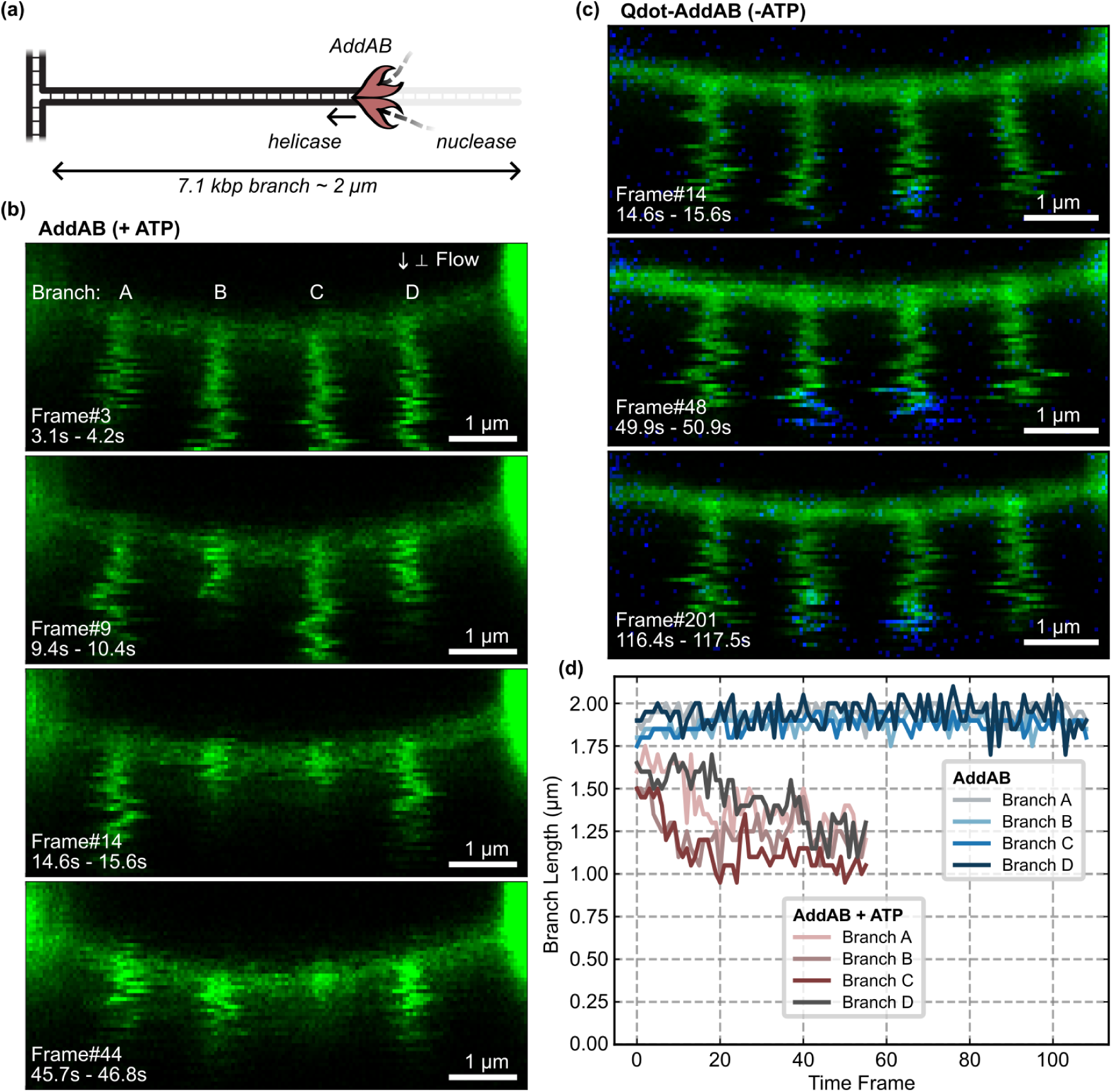
Helicase and nuclease activity of AddAB. **(a**) Upon loading on the DNA end, and in the presence of ATP, helicase/nuclease AddAB can open the duplex while cleaving the nascent strands. **(b)** Time-lapse confocal scans of a long-branched OT-Curtains substrate labelled with SxO in the presence of 10.5 nM AddAB and 250 µM ATP. **(c)** Timelapse confocal scans of the same long-branched DNA shown in (b), labelled with SxO and incubated with 10.5 nM Qdot525-AddAB, in absence of ATP. (**d**) Branch-length trajectories upon exposure to AddAB, in the absence (upper trajectories) and presence (lower trajectories) of ATP.

To quantify branch dynamics, we extracted the apparent branch lengths from the twodimensional confocal fluorescence data (**Fig. 3d**, **Supp. Fig. S3;** see Methods for a detailed description). The branch-backbone junction was defined as the zero point, and the axial DNA baseline established the detection limit, around 15 pixels (750 nm). These trajectories enabled quantification of AddAB activity and rate determination (**Supp. Fig. S4**). For an ATP concentration of 250 µM, one order of magnitude below saturation conditions, AddAB shows a rate of approximately 150 bp/s, a value that agrees with previous reports ^34^. Interestingly, branch shortening often stalled before complete unwinding (**Fig. 3d**). Such stalling, likely reflects the presence of nicks or the generation of a double strand break, either through closely-spaced cleavage events on both strands favoured by the slow translocation (low ATP concentration) or due to a pre-existing nick in one strand. Notably, these apparent stalling events never resumed, suggesting that the DNA-end structures produced by the enzyme were not competent for subsequent AddAB reloading.

Seminal work from the Kowalczykowski laboratory also employed optical tweezers to study the activity of the *E.coli* RecBCD helicase-nuclease by flow-stretching single DNA molecules attached to optically trapped beads and visualizing them by fluorescence microscopy ^20,29,35^. Similar to our approach with AddAB, RecBCD activity was inferred from the shortening of the DNA tether due to the displacement of the dye during unwinding. These pioneering single-molecule studies unveiled the role of Chi sequences and the coordination between helicase motors within the trimeric complex ^35^. However, their measurements were limited to one DNA molecule at a time, making the acquisition of statistically representative data laborious. In contrast, OT-Curtains enables simultaneous tracking of protein-driven unwinding events initiated from several DNA ends, all exposed to identical experimental conditions. The temporal resolution in both approaches is limited by the image acquisition rate. While higher frame rates can be achieved by reducing the scanning area to a single branch (albeit losing the parallelisation), scanning-based methods such as confocal microscopy are inherently slower than full-field imaging methods like TIRF or epi-fluorescence microscopy. The experiments shown in **Fig. 3** were obtained at a rate of 1 s per frame (confocal scanning time across the entire defined area), compared to typical tens-of-milliseconds frame rates attainable with fluorescent cameras.

### Cooperative threading of Ku70-Ku80 onto DNA ends

The Ku heterodimer is a core component of the NHEJ machinery responsible for repairing DSBs in eukaryotic cells. Ku exhibits high affinity for DNA ends and serves as a DSB sensor, as it is thought to be one of the first proteins to bind to broken DNA ends. Once bound, Ku acts as a recruitment platform for other NHEJ factors required to assemble a synaptic complex between the two ends of the break, and to mediate end processing and ligation.

*In vitro* studies have shown that when the DNA fragment is sufficiently long and unoccupied, multiple Ku molecules can cooperatively thread along the DNA ^36,37^. These data suggest that Ku molecules slide inward from the DNA end, exposing the end and allowing the loading of additional Ku units. However, the relevance of this cooperative threading of multiple Ku remains a matter of debate, as measurements in cells estimated that only one or two Ku molecules are loaded per DNA end, and that the rapid association of DNA-PKcs to Ku restricts Ku entry into chromatin ^38^.

To study Ku occupancy at DNA ends using OT-Curtains, we employed a substrate variant containing four short 60 bp branches terminating in blunt ends (**Fig. 4a**; see Methods for details on fabrication). This short-branched configuration ensures a force-free condition at the DNA ends. This is a key feature of our approach because avoids artefacts associated with DNA stretching by hydrodynamic drag, as it is common in flow-stretched methods. The DNA was incubated for 5 min with 50 nM JF549-labelled Ku. During this incubation, Ku heterodimers can thread onto DNA ends and translocate inwards along each branch (**Fig. 4b**). After incubation, the DNA was moved out of the reservoir containing the protein, and a one-dimensional confocal fluorescence scan was recorded along the DNA axis (**Fig. 4c**). 1D-scans considerably improve the time resolution of measurements with typical rates of 20-40 ms per line. Importantly, performing the line scan outside the reservoir improves imaging quality by increasing the signalto-noise ratio.

**Fig. 4.**
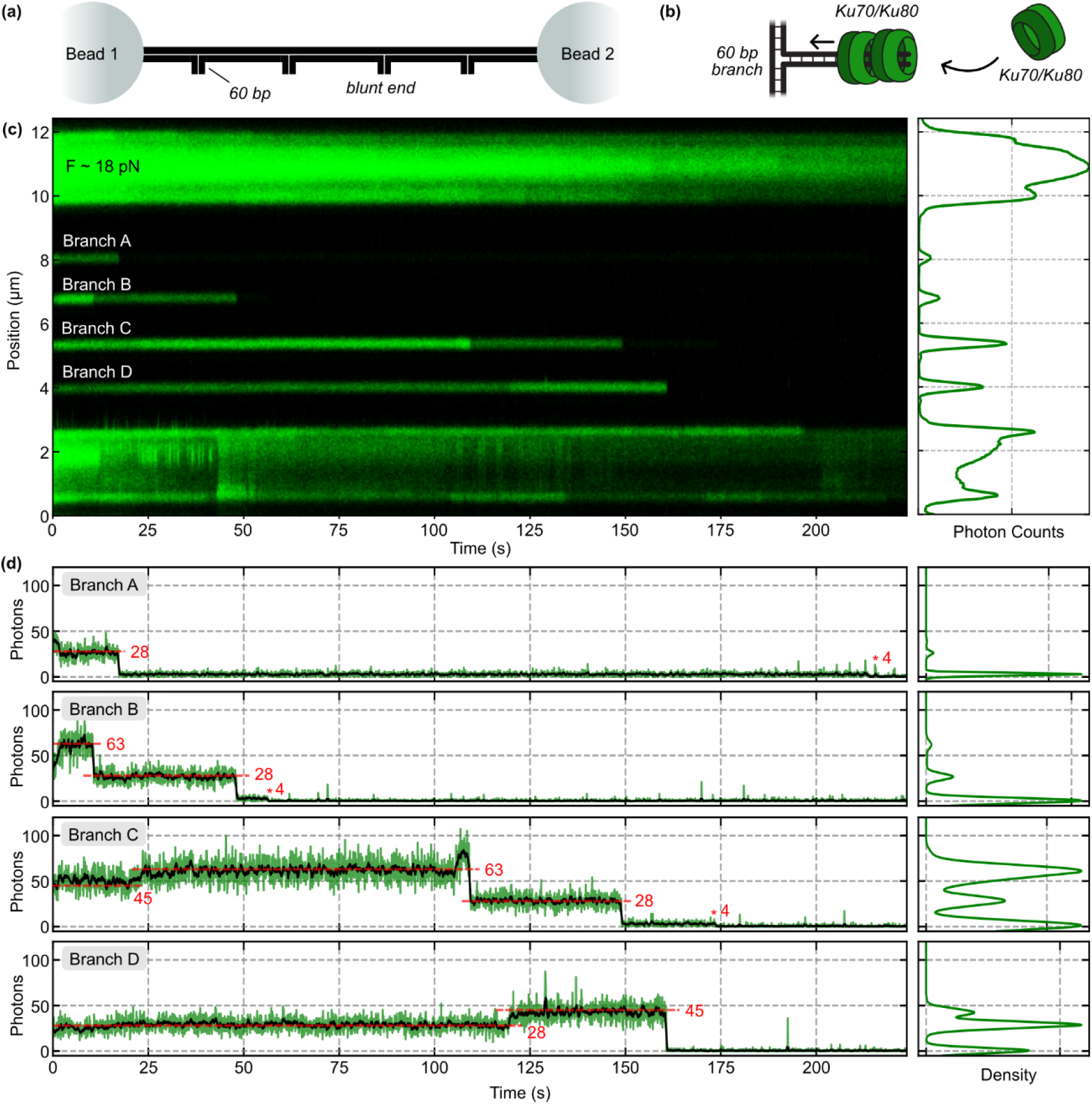
DNA-end specific binding and multiple threading by Ku70-Ku80 heterodimer. **(a)** OT-Curtains with four short (60 bp) branches with blunt ends. **(b)** Detail of a DNA branch with Ku heterodimers sliding inwards. Ku footprint has been reported to be approximately 20 bp. **(c)** Kymograph showing of JF549-labelled Ku binding to DNA ends. Branches display several fluorescent intensity levels. Laser power was increased from 5 % to 15 % to induce the photobleaching of the fluorophores. **(d)** Photon count quantification from an area of 20 pixels of 0.025µm around the individual branches. Four discrete fluorescence intensity levels were detected (63, 45, 28, and 4 photon counts).

By promoting fluorophore photobleaching, we estimated the number of Ku molecules bound to each branch. Photon-count profiles revealed four discrete intensity levels at approximately 63, 45, 28 and 4 photons (**Fig. 4d**). Notably, the three highest photon levels (63, 45, 28 photons) showed intensity differences of roughly 18 photons, suggesting a linear relationship, whereas the lowest level (4 photons) is separated from the next by a 7-fold increase in signal. Fluorescence noise followed a similar trend, being clearly higher for the upper three levels. These observations suggest that the 60 bp branch likely accommodates three Ku heterodimers, in agreement with crystallographic and footprinting studies that estimated a protein coverage of around 20 bp^36,39^, and given that measurement of the labelling ratio of the protein was slightly over one fluorophore per Ku (see methods). The 4 photons level was somehow unexpected, and might reflect some intermediate decay level of this super-bright fluorophore, excited at high power. Further characterization, out of the scope of this work, would be required in this direction.

In a separate assay designed to extend fluorophore lifetime by reducing the laser power to 5 %, the DNA was incubated with JF549-Ku and then transferred to channel 3, where we monitored the dissociation of Ku from the branches. We observed Ku heterodimers remained bound to DNA ends for tens of minutes, with fluorescent signals lasting up to approximately 50 minutes (**Supp. Fig. S5**). This assay is limited by fluorophores’ lifetimes, but it demonstrates the remarkable affinity of Ku to DNA ends and a very low dissociation rate (K_off_).

As shown here, OT-Curtains enables high-resolution studies of protein binding at DNA ends. By labelling the proteins, and intentionally photobleaching the fluorophores, we can quantify the number of molecules bound to each end. This configuration is useful not only for analysing DNA-end binding, but also for investigating processes catalysed at DNA ends. For example, by reconstituting the NHEJ core machinery at the DNA ends and introducing fluorescently labelled DNA hairpins into the solution, one could monitor the formation of DNA end synapsis under varying conditions or in the presence of different proteins and/or RNA factors. Similarly, this platform could also be applied to study the HR machinery. The resection step could be examined by reconstituting the relevant complexes and following the progression of resection through changes in fluorescence signal.

### DNA condensation by ParB upon *parS* docking

ParB is a CTP-dependent DNA-binding protein that organizes bacterial chromosomes as part of the ParABS partitioning system ^40,41^. It nucleates at *parS* sites, typically clustered near the origin of replication, where sequence-specific binding promotes CTP binding and converts ParB into a closed, sliding clamp ^40–42^. Once loaded at *parS*, ParB spreads and diffuses along adjacent dsDNA, forming a dynamic nucleoprotein zone capable of bridging DNA segments and condensing the locus ^42–44^. CTP hydrolysis promotes clamp opening and protein turnover, enabling cycles of loading at *parS* and release from flanking DNA ^45^. Through this CTP-driven assembly-disassembly cycle, ParB establishes the partition complex, recruits and modulates SMC condensin activity at the origin, and interacts with the ParA ATPase to coordinate chromosomes position and segregation ^46–48^.

The OT-Curtains approach is also suitable for studying processes that do not require DNA ends but benefit from the force-free configuration and the freedom of movement that OT-Curtains provide. Here, we employed a variant containing four long 6.2 kbp branches terminating in blunt ends and harboring a single *parS* site (**Fig. 5a**; see Methods for details on fabrication). When the branches were flow-stretched in the presence of 200 nM ParB dimer and 1 mM CTP, we observed progressive condensation of the long branches (**Fig. 5b, c**, and **Supp. Video 4**). After approximately 1 minute, the 6.2 kbp branches were fully condensed. A limitation of OT-Curtains is the minimum detectable branch length. Branches shorter than 0.7 µm (14 pixels at 0.05 µm per pixel) cannot be resolved, as their fluorescence signal merges with that of the DNA backbone (**Fig. 5c**, **Supp. Fig. S3**). For a 6.2 kbp branch with an initial extension of 1.7 µm, the detection limit corresponds to approximately 2.5 kbp; shorter extensions cannot be distinguished.

**Fig. 5.**
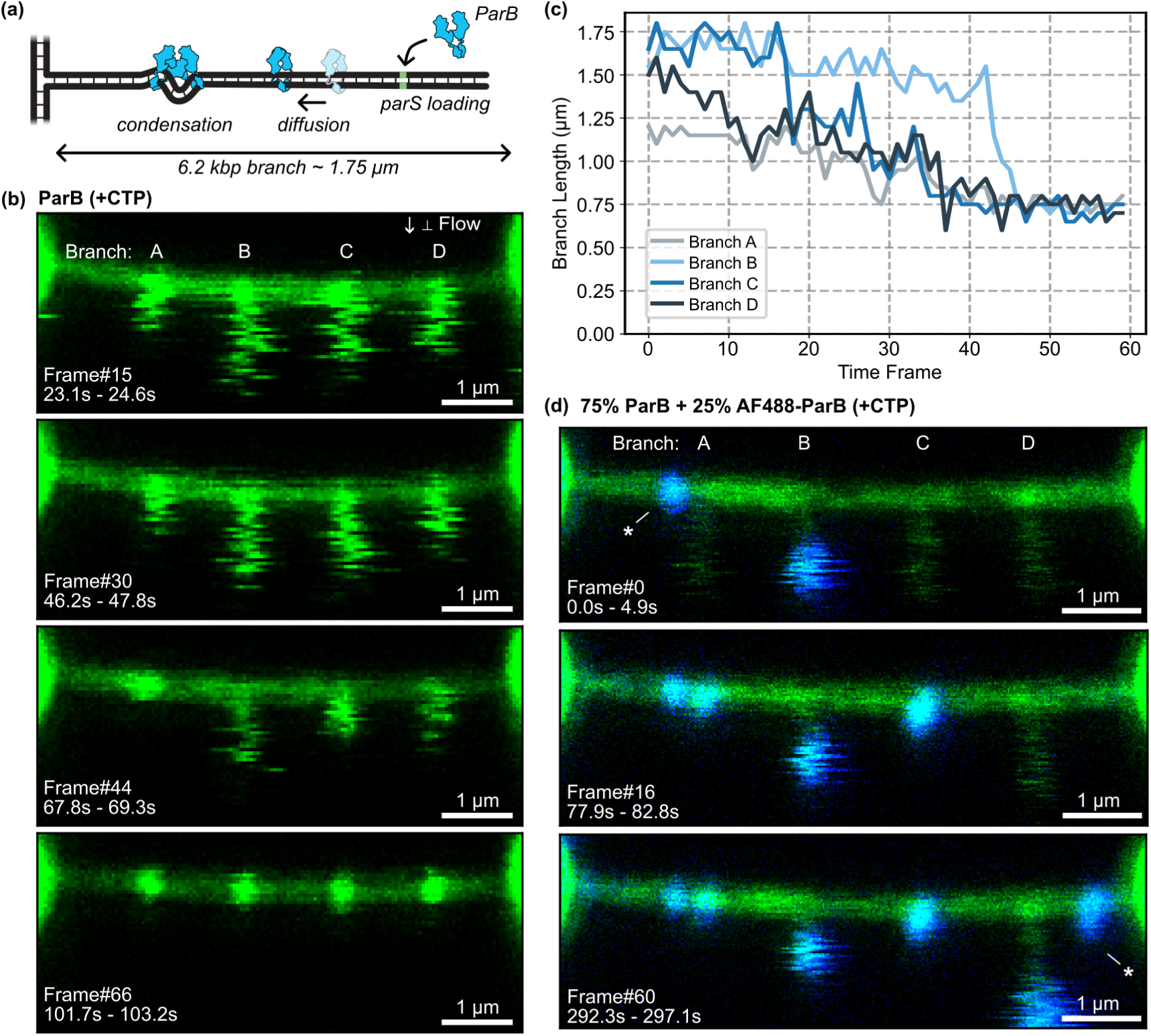
DNA condensation by ParB. **(a)** Upon loading on a parS site, ParB diffuses along the branch and interacts with other previously loaded ParB molecules to create bridges on the DNA. **(b)** Time-lapse confocal scans of OT-Curtains with 6.2 kbp long-branches containing a single parS sequence in the presence of 200 nM ParB dimer and 1 mM CTP, and 100 nM SxO to visualize the DNA. **(c)** Branch-length trajectories for ParB-mediated DNA condensation. **(d)** Introducing a small percentage of labelled ParB (Alexa Fluor 488-ParB) allows us to monitor the binding of individual molecules. Sites marked with an asterisk (*), represent parS sites on the DNA backbone.

To monitor individual ParB molecules during DNA condensation, we repeated the OT-Curtains assay in the presence of 75 % unlabelled ParB and 25 % Alexa Fluor 488-labelled ParB. Results showed ParB binding to DNA at different stages of the condensation (**Fig. 5d, Supp. Video 5**). During the initial stages it seemed to bind close to the end of the branches (**Fig. 5d**, Branch A and D), where the single *parS* site is located. In later stages, the signal is observed on the fully condensed branch (**Fig. 5d**, Branch C). In addition, ParB was also spotted on the *parS* sites located on the backbone DNA, which is tensioned, in a configuration incompatible with condensation. (**Fig. 5d**, asterisk).

Rapid buffer exchange enables exploration of condensation dynamics under varying conditions. ParB condensation of DNA molecules containing a single *parS* site has proven to be challenging in MT experiments, where even very low pulling forces prevented DNA condensation, and substrates containing several *parS* sites were employed ^44^. However, alternative methods aimed at minimizing the force acting on the DNA successfully captured DNA condensation by parB on single *parS* substrates^49^. Here we show DNA condensation by ParB on substrates with only one *parS* site, demonstrating that the flow used to extend the DNA applies a very low force which remains compatible with DNA condensation by ParB proteins. Weak condensation activities as those exhibited by the SMC5/6 complex are also amenable to be studied using OT-Curtains^50^.

As shown here, OT-Curtains enables monitoring of condensation dynamics without imposing mechanical constraints on the molecular components. By introducing specific DNA sequences and adjusting buffer conditions, OT-Curtains can be used to systematically examine how varying factors influence condensation kinetics and stability.

### DNA digestion by the restriction enzyme KpnI

The OT-Curtains approach enables the study of restriction enzymes activity without loss of the DNA tether. Restriction enzymes are central to both bacterial physiology and modern biotechnology. In bacteria, restriction endonucleases act with cognate methyltransferases in restriction-modification systems to protect the host from invading DNA, and these systems shape host-phage co-evolution and genome architecture ^51–53^. In biotechnology, their sequencespecific cleavage activity serves as a fundamental tool for molecular cloning, library construction, DNA barcoding, and mapping. Mechanistic insights into how restriction enzymes recognize target motifs and navigate obstacles provide valuable information for enzyme engineering, off-target analysis, and the rational design of DNA substrates^54^. Understanding these mechanisms enhances experimental control over DNA cleavage and may yield new insights into native cellular surveillance processes.

To demonstrate the versatility of OT-Curtains for studying restriction enzymes, we examined the endonuclease activity of KpnI, a commercially available enzyme that recognizes and cleaves the 5′-GGTAC/C-3’ sequence. We employed an OT-Curtains variant containing four long 7 kbp branches terminating in blunt ends (see Methods for details on fabrication). Each branch included a KpnI recognition site located 1.8 kbp from the backbone-branch junction.

Exposure of the OT-Curtains substrate to a high concentration of KpnI resulted in branch cleavage as expected (**Fig. 6a, b**). Notice that during the experiment, branches C and D exhibited transient abnormal motion lasting several frames (see **Fig. 6b**, frames 25-30, and **Supp. Video 6**). This motion was detected in the branch-length trajectories, highlighting the sensitivity of the measurement. Cleavage at the KpnI site releases a 5.3 kbp fragment (**Fig. 6c**). Quantification of the branch lengths after digestion showed that KpnI cleaved at the predicted site (90% of events) but eventually also revealed two additional non-specific nearby cuts (2/20 events) (**Fig. 6d**). This might be the result of unspecific photocleavage.

**Fig. 6.**
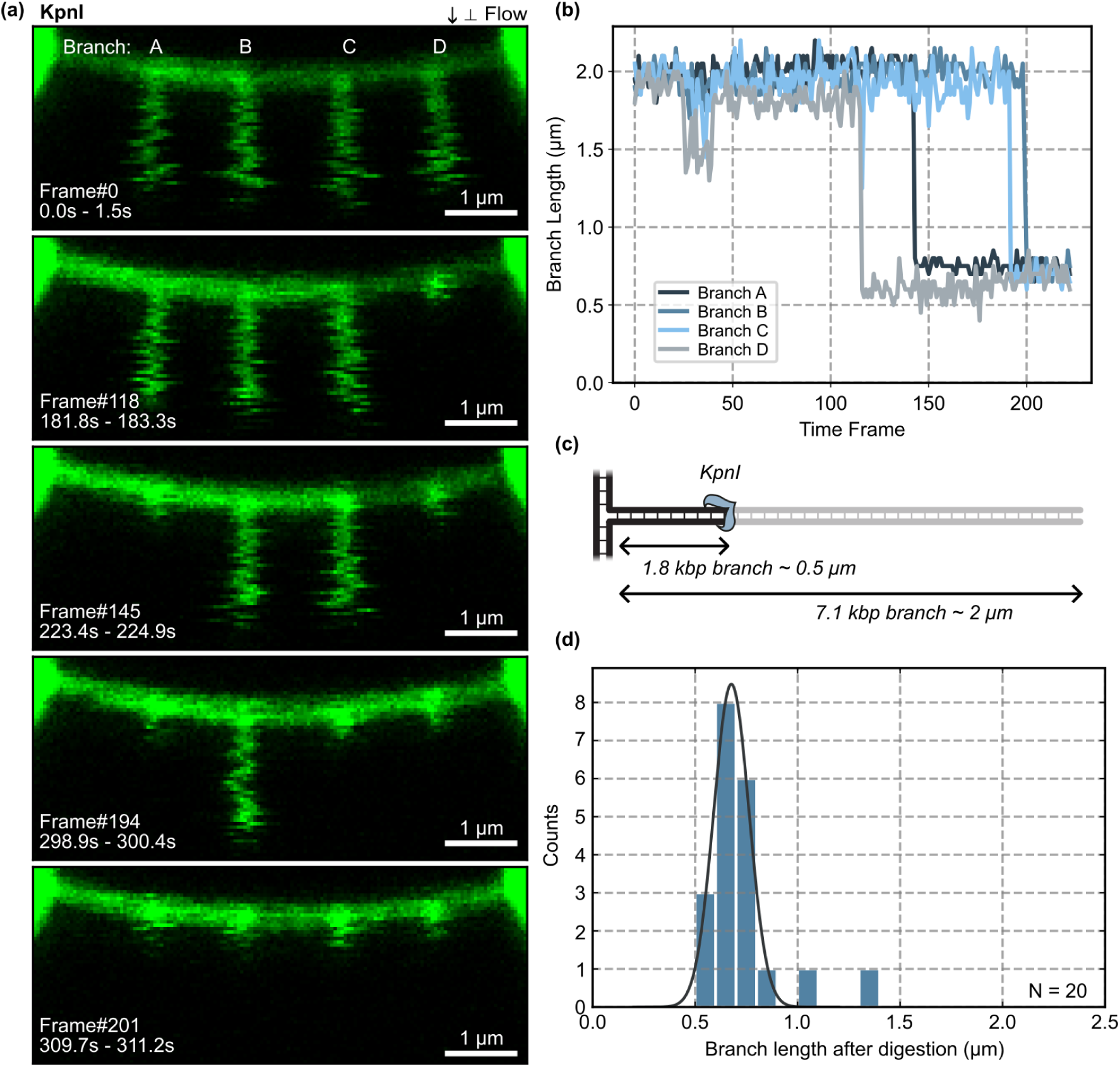
DNA cleavage by KpnI restriction enzyme. **(a)** Time-lapse confocal scans of OT-Curtains with 7 kbp long-branches containing a single KpnI recognition site, showing progressive DNA cleavage over time. **(b)** Branch-length trajectories in the presence of KpnI. **(c-d)** Schematics and experimental distribution of branch length after digestion by KpnI. N indicates the number of cleaved branches.

As shown here, the OT-Curtains enables real-time monitoring of sequence-specific cleavage without loss of the tethered DNA molecule. This approach provides a means to systematically evaluate restriction enzyme specificity and reaction conditions in a force-free, single-molecule context.

## Conclusions

In this work, we introduce OT-Curtains, a platform for single-molecule studies of DNA-protein interactions at accessible DNA ends. The method employs dual-trap optical tweezers to hold a custom DNA construct from which several branches emerge, and integrated confocal microscopy for visualization of fluorescently labelled DNA molecules and proteins. OT-Curtains enables simultaneous, force-controlled observation of end-specific processes such as DNA end binding and resection, as well as other reactions that do not necessarily require a DNA end, including DNA condensation and restriction enzyme activity. All these applications benefit from the exceptionally low forces applied to the DNA, the multiplexing capacity of the assay, the structural integrity of the substrate upon cleavage on the branches, and the rapid exchange of reagents that facilitates the exploration of diverse experimental conditions.

The versatility of OT-Curtains was demonstrated across multiple biological systems, including AddAB helicase–nuclease activity, Ku70–Ku80 loading, ParB-mediated DNA condensation, and KpnI restriction endonuclease activity. Among these, the most impactful applications lie in the field of DNA double-strand break repair and processing, where OT-Curtains provides a unique opportunity to visualize and quantify DNA-end transactions in real time. For this application, the short branch configuration is particularly powerful because protein-induced branch activity can be captured with rapid line scans along the OT-Curtains backbone of tens of tens of milliseconds, achieving the highest temporal resolution of all configurations.

OT-Curtains enables direct observation of fluorescently-labelled proteins binding to a variety of DNA end structures, including forks, tails, hairpins, and modified oligonucleotides. The subsequent fate of these bound proteins, such as their removal or remodelling by other factors or multiprotein assemblies, can be investigated within the same experimental framework. This capability opens the door to addressing central questions in DNA repair, for instance: What is the molecular mechanism behind the removal of DNA-end-bound Ku proteins by the HR machinery? Or, how do repair factors and RPA coordinate during DNA-end processing?

The current OT-Curtains design employs a backbone containing four identical ssDNA gaps; however, further engineering could increase the number of BbvCI sites, thereby enhancing multiplexing capacity and including different gap sequences. The spatial separation of these gaps is limited by the diffraction limit of confocal fluorescence microscopy (λ/2NA, typically a few hundred nanometers). Nonetheless, integration with super-resolution fluorescence microscopy could overcome this restriction. In addition, tandem constructs can be optically trapped to double the number of branches, ssDNA gaps, or DNA ends (**Supp. Fig. S1**). This can be easily achieved by adjusting the ratio of backbone to handles during ligation. The possibility to incorporate different types of branches along the backbone enables the simultaneous examination of sequence-dependent effects on protein binding, including internal controls, within a single experiment.

Finally, the backbone DNA, which forms the cornerstone of this platform, offers broad utility for studying the binding dynamics of single stranded DNA-binding proteins and their interaction with other DNA-remodeling factors. RPA and SSB proteins are central to virtually all DNA transactions that generate ssDNA in eukaryotes and bacteria, respectively. The ssDNA-gapped OT-Curtains backbone provides an ideal substrate to monitor their binding, as well as the production of additional ssDNA by helicases or exonucleases involved in HR. Moreover, the system is well suited to investigate their replacement by recombinases RAD51 in eukaryotes and RecA in bacteria, using the backbone DNA containing multiple ssDNA gaps.

In summary, OT-Curtains provides a versatile tool for probing DNA-protein interactions by addressing two challenges that have traditionally been inaccessible to optical tweezers users: multiplexing and accessible DNA ends. OT-Curtains overcomes these long-standing limitations and opens the possibility for investigating molecular motors, repair factors, and complex protein assemblies acting on DNA in ways that were previously unattainable.

## Online content

Full protocols, descriptions and specific parameters, as well as additional references, are provided in the Online Methods. Funding sources are listed in Acknowledgements. Author roles are detailed in Author contributions. Data and code are available at a Zenodo repository ^55^ and Moreno-HerreroLab Github repository at https://github.com/Moreno-HerreroLab/CTrapDataProcessing.

## Online Methods

### Fabrication of a large plasmid with four poly-BbvCI regions

Experiments were performed using a linear double-stranded DNA (dsDNA) molecule of approximately 24 kbp containing four dsDNA branches, either long or short. To generate these DNA molecules, we built a large plasmid that incorporated four distinct regions, each composed of five closely spaced BbvCI restriction sites (poly-BbvCI regions), as described below.

An initial plasmid containing one poly-BbvCI region (*1x poly-BbvCI* plasmid), first reported in Aicart-Ramos *et al.* ^56^, was obtained by extracting a 1,428 bp fragment carrying the poly-BbvCI region from the plasmid reported in Wilkinson *et al.* ^57^, originally extracted from the pNLrep plasmid ^58^ and by inserting itinto the XhoI site of the plasmid *C-trap EcoRI 39 parS* reported in Balaguer *et al.* ^59^. The result is a 18,772 bp plasmid containing one poly-BbvCI region.

To introduce a second poly-BbvCI region, the same 1,428 bp fragment was inserted into de SalI site of the *1x poly-BbvCI* plasmid. For this, the plasmid was digested with SalI (New England Biolabs, #R0138S), dephosphorylated using rSAP (New England Biolabs, #M0371S), and without further purification, ligated with the fragment using T4 DNA ligase (New England Biolabs, #M0202S). *E. coli* DH5α competent cells (Thermo Fisher Scientific, #18265-017) were transformed with the ligation product, and potentially positive colonies were screened by colony PCR. Plasmids were purified using the QIAprep Spin Miniprep Kit (QIAGEN, #27106), verified by restriction enzyme digestion, and finally confirmed by DNA sequencing. This resulted in a 20,200 bp plasmid containing two poly-BbvCI regions separated by 6,459 bp (*2x poly-BbvCI* plasmid).

For the third poly-BbvCI region, two complementary long oligonucleotides carrying the poly-BbvCI cassette (**Supplementary Table S2, Frag. A**) were annealed by heating to 95 °C for 5 min and cooling to 20 °C at -1 °C min⁻¹ in 10 mM Tris-HCl (pH 8.0), 1 mM EDTA, 200 mM NaCl, and 5 mM MgCl₂. The duplex was phosphorylated at the 5’ termini using T4 polynucleotide kinase (New England Biolabs, #M0201S) and inserted into the PciI site of the 2x poly-BbvCI plasmid. The recipient plasmid was digested with PciI (New England Biolabs, #R0655S), dephosphorylated with rSAP, and ligated with the phosphorylated duplex using T4 DNA ligase. Following transformation and screening as above, a 20,278 bp plasmid containing three poly-BbvCI regions in the same orientation was obtained (*3x poly-BbvCI* plasmid).

To introduce the fourth poly-BbvCI region, the *p64.large* plasmid backbone described by Sukhoverkov *et al.* ^60^ was used as template to amplify a new dsDNA fragment containing a poly-BbvCI region by PCR using Phusion High-Fidelity DNA Polymerase (Thermo Fisher Scientific, #F553S). The primers introduced BstBI restriction sites at both ends (see **Supplementary Table S2, Frag. B**). After purification using the QIAquick PCR Purification Kit (QIAGEN, #28106) followed by BstBI digestion (New England Biolabs, #R0519S), the fragment was purified by gel extraction using the QIAquick Gel Extraction Kit (QIAGEN, #28706). The plasmid containing three poly-BbvCI regions was digested with BstBI, dephosphorylated with rSAP, and ligated to the purified fragment using T4 DNA ligase. The new cassette was inserted between the first and second poly-BbvCI regions (originally separated by 6,459 bp). Colonies were screened and sequence-verified as described above, resulting in the final 22,142 bp plasmid containing four poly-BbvCI regions (*4x poly-BbvCI* plasmid), which was used to prepare the OT-Curtains dsDNA constructs (**Fig. 1a**)

### DNA construct with four single-stranded DNA gaps

The backbone of the OT-Curtains construct is a linear double-stranded DNA (dsDNA) molecule of 22,138 bp with 4-nucleotide 5’ cohesive ends. It contains four 63-nucleotide single-stranded DNA (ssDNA) gaps separated by 3,693 bp, 4,211 bp, and 3,986 bp, and positioned 5,571 bp and 4,425 bp from the DNA termini, respectively (**Fig. 1c**).

The construct is prepared as follows. The 22,142 bp *4x poly-BbvCI* plasmid (see previous section) is digested with BssHII (New England Biolabs, #R0199S) to linearize the DNA. Simultaneously, the plasmid is treated with the nicking enzyme Nt.BbvCI (New England Biolabs, #R0362S). Digestion of each poly-BbvCI region by Nt.BbvCI introduces five nicks on the same DNA strand (**Fig. 1b**). During subsequent heat inactivation of the enzymes, the short 15-16 nt fragments generated by nicking are released, resulting in the formation of four 63-nucleotide singlestranded gaps (**Fig. 1c**).

The resulting linear dsDNA with four ssDNA gaps (**Supplementary Table S3**) is used directly, without further purification, as the backbone molecule for OT-Curtains assembly. These ssDNA gaps serve as hybridization sites that can be filled with complementary oligonucleotides to create defined structures, including ssDNA flaps, dsDNA branches, or more complex DNA architectures, depending on the biological application ^57,58,61,62^.

### DNA handles, biotinylated dsDNA fragments for DNA immobilization

Biotinylated dsDNA fragments are required at both ends of the OT-Curtains constructs to enable optical trapping and molecular immobilization; these fragments are referred to as handles.

A ∼1 kbp-long, highly biotinylated DNA fragment was prepared by PCR amplification (see **Supplementary Table S2, Frag. C** for primers) using GoTaq® G2 Flexi DNA Polymerase (Promega, #M7805). The PCR reaction contained 200 µM each of dGTP, dCTP, and dATP, along with 140 µM dTTP (Roche, #11 969 064 001) and 66 µM Bio-16-dUTP (Roche, #11 093 070 910), using the plasmid pSP73-JY0 ^63^ as the template, following the procedure described in ^56^.

To generate cohesive ends compatible with the OT-Curtains backbone, the PCR amplicons were purified using the QIAquick PCR Purification Kit, then digested with MluI-HF (New England Biolabs, #R3198S) and again purified using the QIAquick PCR Purification Kit. The resulting biotinylated fragment was used as the DNA handle for all OT-Curtains construct variants described in this work.

### OT-Curtains with long ATTO488-labelled branches (7 kbp)

To fabricate the OT-Curtains substrate with 7 kbp-long branches labelled with ATTO488 at position 33 from the branch end, the DNA construct with four ssDNA gaps was first prepared as described in a previous section. The ssDNA gaps were then filled with the long-branched structures (**Fig. 2a**). The sequence of the long branch is provided in **Supplementary Table S4**.

The long-branched structures were assembled by annealing and ligating a small connector to a PCR-amplified DNA fragment. The connector consisted of a 40 bp duplex stem terminating at one end in a 12 nt 3’ overhang complementary to the 3’ overhang of the long PCR amplicon, and at the opposite end carrying two single-stranded arms of 31 and 32 nts complementary to the 63 nt ssDNA gaps generated at the poly-BbvCI regions. The connector was prepared by annealing two partially complementary oligonucleotides (**Supplementary Table S2, Frag. G**) by heating to 95 °C for 5 min, followed by slow cooling to 20 °C at -1 °C min⁻¹ in 10 mM Tris-HCl (pH 8.0), 1 mM EDTA, 200 mM NaCl, and 5 mM MgCl₂.

The long PCR fragment (7,108 bp) was amplified using Phusion High-Fidelity DNA Polymerase (Thermo Fisher Scientific) and the pSP73-JY0 plasmid ^63^ as template (see **Supplementary Table S2, Frag. E** for primers). The amplicon contained a BbvCI restriction site at one end and was purified using the QIAquick PCR Purification Kit. To create a 12 nt 3’ overhang, the purified fragment was digested with the nicking enzyme Nt.BbvCI, following established protocols ^58,64^.

The 12 nt 3’ overhang of the PCR fragment was annealed to the connector using a 25-fold molar excess of the latter. The mixture was heated to 72 °C for 10 min and slowly cooled to 42 °C at a rate of -0.1 °C every 25 s in 10 mM Tris-HCl (pH 7.5) and 1 mM MgCl₂. The annealed fragments were ligated overnight at 16 °C using T4 DNA Ligase. The resulting long-branched DNA structures (7,148 bp) were gel-extracted and purified.

A 20-fold molar excess of these long-branched structures was then hybridized into the linear dsDNA backbone containing four ssDNA gaps. The mixture was heated to 80 °C for 5 min and cooled to 30 °C at -0.5 °C min⁻¹, increasing the dwell time by 10 s per minute, in 50 mM Tris-HCl (pH 8.0), 1 mM EDTA, and 100 mM NaCl. Finally, biotinylated handles (prepared as described previously) were ligated to the linear OT-Curtains at a 20-fold molar excess of handles to backbone using T4 DNA Ligase for 15 h at 16 °C. This ligation step also sealed all remaining nicks.

The resulting OT-Curtains constructs with 7 kbp-long ATTO488-labelled branches were ready for use without further purification. Constructs containing eight branches could be obtained by adjusting the ratio of handles to dsDNA backbones during ligation, thereby generating tandem molecules. The length and sequence of the branches can be customized for specific applications; however, branch fragments must not contain internal BbvCI restriction sites.

### OT-Curtains with short ATTO488-labelled branches (20 bp)

To fabricate OT-Curtains constructs with 20 bp-long short branches labelled with ATTO488 at the branch termini (**Fig. 2c**), the DNA backbone containing four ssDNA gaps was first prepared as described above.

Short branches were assembled as in de Bragança *et al.* ^62^ by annealing two partially complementary oligonucleotides to form a 20 bp duplex stem with a blunt end on one side and two non-complementary single-stranded extensions on the opposite side. These 31 and 32 nt-long extensions were designed to be complementary to the 63 nt ssDNA gaps generated at the poly-BbvCI regions, thereby fully filling them upon annealing.

Oligonucleotides for the short branches (see **Supplementary Table S2, Frag. D**) were annealed by heating to 95 °C for 5 min and cooling to 20 °C at -1 °C min⁻¹ in 10 mM Tris-HCl (pH 8.0), 1 mM EDTA, 200 mM NaCl, and 5 mM MgCl₂. The resulting short-branch structures were then hybridized to the OT-Curtains backbone at a 200-fold molar excess. The mixture was heated to 80 °C for 5 min and cooled to 30 °C at -0.5 °C min⁻¹, increasing the dwell time by 10 s per minute, in 50 mM Tris-HCl (pH 8.0), 1 mM EDTA, and 100 mM NaCl, as described previously ^57^.

Finally, biotinylated handles (prepared as described above) were ligated to the OT-Curtains backbone at a 20-fold molar excess of handles to backbone using T4 DNA Ligase for 15 h at 16 °C. This ligation step sealed all remaining nicks.

The resulting short-branched OT-Curtains constructs required no further purification and were ready for use. Constructs bearing eight branches could be obtained by reducing the ratio of handles to backbone during ligation, promoting tandem association of the backbone molecules. Branch length, sequence, and fluorophore labelling can be readily customized depending on the specific application.

### OT-Curtains with long (unlabelled) branches (7 kbp)

To fabricate OT-Curtains constructs with 7 kbp-long branches (used in **Fig. 3**), we followed the same procedure described for the preparation of OT-Curtains with 7 kbp-long ATTO488-labelled branches, with the exception of the PCR fragment used.

In this case, the PCR amplicon was generated using the pSP73-JY0 plasmid as a template but employing a different set of primers (see **Supplementary Table S2, Frag. H**). The sequence of the 7 kbp-long branches used in these constructs is provided in **Supplementary Table S5**.

### OT-Curtains with short (unlabelled) branches (60 bp)

To fabricate OT-Curtains constructs with 60 bp-long short branches (**Fig. 4**), we followed the same procedure described for the preparation of OT-Curtains with short ATTO488-labelled branches, using a different pair of partially complementary oligonucleotides.

In this case, two oligonucleotides (see **Supplementary Table S2, Frag. F**) were annealed to form a 60 bp duplex stem with a blunt end, replacing the shorter fluorescently labelled counterparts used previously.

### OT-Curtains with 6.2 kbp-long branches containing one *parS* site

To fabricate OT-Curtains constructs with 6.2 kbp-long branches containing a single *parS* site (**Fig. 5**), we followed the same procedure described for the preparation of OT-Curtains with 7 kbplong ATTO488-labelled branches, using a different PCR-amplified fragment.

In this case, the fragment was amplified from the plasmid reported by Taylor *et al.* ^65^ using the primers listed in **Supplementary Table S2, Frag. I**. This plasmid contains the optimal *Bacillus subtilis parS* sequence (5’-TGTTCCACGTGAAACA-3’). The complete sequence of the 6.2 kbp-long branch containing one *parS* site is provided in **Supplementary Table S6**.

### Fluorescent labelling of Ku

The Ku heterodimer used for fluorescent labelling experiments (Halo-Ku) consisted of full-length Ku70 fused at its N-terminus to a Twin-Strep tag followed by a Halo tag, and full-length Ku80 carrying an N-terminal 10×His tag. The construct was expressed and purified as described previously ^62,66^.

The Janelia Fluor® Dye JF549i-HaloTag ligand was kindly provided through the Open Science initiative by Janelia Research Campus. The dye was dissolved in dimethyl sulfoxide (DMSO; Sigma) to a final concentration of 1 mM.

Labelling reactions were performed in a total volume of 220 µL containing 34 µM Halo-Ku, 80 µM dye, 1 mM DTT, and 25 mM HEPES-NaOH (pH 7.0). The mixture was incubated overnight at 4 °C with gentle rocking and protected from light.

Labelled protein was separated from unreacted dye by size-exclusion chromatography on a Superdex 200 Increase 10/300 GL column (Cytiva, #GE17-5175-01) equilibrated in 25 mM HEPES-NaOH (pH 7.0), 250 mM NaCl, 5% (w/v) glycerol, and 0.5 mM TCEP. Peak fractions were monitored by absorbance at 280 nm, 549 nm, and 571 nm. Apparent labelling efficiency was assessed by spectrophotometry as described by the vendor, by comparing the absorbance from the protein and the JF549i dye, including a correction factor for absorbance of the dye at 280 nm (0.169). Ku was completely labelled, as we calculate a degree of labelling of 1.1 moles of dye per mole of Ku heterodimer.

### Confocal fluorescence microscopy-correlated Optical Tweezers (C-Trap) assays

Correlative trapping-fluorescence experiments were performed at room temperature using a commercially available integrated system that combines dual-trap optical tweezers with three-color confocal fluorescence microscopy and computer-controlled stage that enables rapid positioning of the optical traps within a five-channel microfluidic flow cell (C-Trap; LUMICKS).

The microfluidic chamber is designed to feature five different medium channels separated by laminar flow. Channels 1 to 3 were used as main channels that allow for bead trapping and DNA immobilization. Channels 4 and 5, were used as reservoirs for the proteins of study. The chamber outlet is referred to as channel 6.

The instrument is equipped with three excitation lasers (488 nm, 532 nm, and 639 nm) enabling multi-color confocal imaging. Emission is collected using a 500-525 nm (blue), a 545-620 nm (green) and a 650-750 nm (red) filters, corresponding to the respective excitation wavelengths.

Kymographs (1D) were generated by scanning across the trapped beads and DNA tethers, whereas 2D confocal scans were acquired over user-defined regions of interest (ROI). The Ztelescope position was adjusted to 3.74 µm to align the confocal focal plane to the tethered DNA molecule.

### Bead trapping and DNA immobilization

The buffer used in the main microfluidic channels (channels 1-3) contained 20 mM HEPES-KOH (pH 7.8), 100 mM KCl, and 5 mM MgCl₂, hereafter referred to as Main Buffer (MBuffer). For all assays described in this work, DNA tethering between two beads followed the same experimental workflow.

In channel 1, a 1:1000 dilution of 1.76 µm streptavidin-coated polystyrene beads (Spherotech, Inc., SVP-15-5) in MBuffer was introduced. When trapping beads for the first time, the optical traps were calibrated, and the laser power was adjusted to achieve a trap stiffness of ∼0.4 pN·nm⁻¹ in each trap.

Then, trapped beads were exposed to channel 2, where a 6:300 dilution of the appropriate dsDNA construct (or 12:300 for constructs containing long branches) in MBuffer was introduced. DNA tethering between the two trapped beads was verified by displacing trap 2 relative to trap 1 and monitoring the resulting force-extension response.

Channel 3 contained MBuffer supplemented with 1 mM DTT and 1 mM Trolox (Sigma-Aldrich, #238813-1G) and was used for imaging and manipulation after tether formation.

### Proof-of-principle assays

Bead trapping and DNA immobilization were performed as detailed in the previous section.

For the proof-of-principle assay with long branches, we used the OT-Curtains with long ATTO488-labelled branches (7 kbp) and the 488 nm-wavelength laser at 5% power. The experimental workflow proceeded as follows. First, a DNA molecule was tethered between the two beads. With all valves closed and vented, the tethered DNA molecule was positioned at a reservoir channel (channel 4 or 5) containing 20 mM HEPES-KOH (pH 7.8), 100 mM KCl, 5 mM MgCl₂, 1 mM DTT, and 1 mM Trolox. The applied force on the DNA was adjusted to 20 pN and the system pressure was set to 0.8 bar. A 2D confocal scan was then initiated, and immediately, channels 6 and selected reservoir channel were opened.

For the proof-of-principle assay with short branches, we used the OT-Curtains with short ATTO488-labelled branches (20 bp) and the lasers with wavelengths 488 nm and 561 nm at 5% and 10% power, respectively. The experimental workflow proceeded as follows. First, a DNA molecule was tethered between the two beads. Then, the tethered molecule was transferred into a reservoir channel (channel 4 or 5) containing 20 mM HEPES-KOH (pH 7.8), 100 mM KCl, 5 mM MgCl₂, 1 mM DTT, and 1 mM Trolox. A 1D kymograph scan was then initiated along the DNA axis.

### AddAB unwinding and cleavage experiments

The biotinylated *Bacillus subtilis* AddAB complex (bio-AddAB) used in these assays was previously described in Carrasco *et al.* ^67^. Bead trapping and DNA immobilization were performed as detailed in the previous section. For these assays, the 532 nm excitation laser was set to 5% power.

The experimental workflow proceeded as follows. First, a DNA molecule was tethered between the two optically trapped beads. With all valves initially closed and vented, the tethered DNA molecule was positioned at the junction between channel 3 and the protein reservoir channel. The applied force on the DNA was adjusted to sufficiently stretch the axial DNA molecule (usually 20 pN), and the system pressure was set to 0.8 bar, while keeping all valves closed. A 2D confocal scan was then initiated. Immediately, channels 6 (output) and protein reservoir channel (channel 4 or 5) were opened, followed by the transfer of the DNA molecule into the selected reservoir.

The protein reservoir contained 10 nM bio-AddAB in 25 mM Tris-OAc (pH 7.5), 2 mM MgOAc, 1 mM DTT, 1 mM Trolox, 100 nM Sytox Orange, and 250 µM ATP (when present). In some experiments, bio-AddAB was pre-incubated with streptavidin-conjugated Qdot525 (Thermo Fisher Scientific, Q10143MP) at a 1:2 (AddAB:Qdot) molar ratio for 10 min on ice before dilution with experimental buffer to a final protein concentration of 10 nM (referred to as QD525AddAB).

### Ku70/Ku80 DNA-end binding experiments

The JF549i-labelled Halo-Ku70/Ku80 heterodimer (JF549-Ku) used in these assays was prepared as described in a previous section. Bead trapping and DNA immobilization were performed as outlined above.

Once the DNA molecule was tethered between the two beads, the 532 nm excitation laser was set to 5% power. The tethered DNA was moved into the designated protein reservoir (channel 4 or 5), which contained 50 nM JF549-Ku in 20 mM HEPES-KOH (pH 7.8), 100 mM KCl, 5 mM MgCl₂, 1 mM DTT, and 1 mM Trolox After a 5-min incubation, the tethered DNA molecule was transferred to the cross junction between Channel 3 and the selected reservoir and the 1D kymograph acquisition was then initiated.

### ParB condensation experiments

The *Bacillus subtilis* ParB and the Alexa Fluor 488-labelled *Bacillus subtilis* ParB (AF488-ParB) proteins used in these assays were previously described in Balaguer *et al.*^59^. Bead trapping and DNA immobilization were performed as outlined in the previous section.

Once the DNA molecule was tethered between the two beads, the 488 nm excitation laser was set to 5% power. The workflow for these experiments was as follows. With all valves initially closed and vented, the tethered DNA molecule was positioned at the junction between channel 3 and the protein reservoir channel (channel 4 or 5). The applied force on the DNA was adjusted to 20 pN, and the pressure was set to 0.8 bar while keeping all valves closed. A 2D confocal scan was then initiated, followed by opening of channel 6 and the relevant reservoir channel. Immediately, the DNA tether is moved into the selected reservoir.

For these assays, the reservoir contained ParB, either unlabelled or AF488-labelled, in 40 mM Tris-HCl (pH 7.5), 65 mM KCl, 2.5 mM MgCl₂, 1 mM DTT, 1 mM Trolox, 1 mM CTP, and 100 nM Sytox Orange.

### KpnI restriction experiments

The KpnI enzyme used in this work was a non-high-fidelity commercial version that has since been discontinued (New England Biolabs, #R0142; discontinued in 2021). This enzyme preparation exhibits star activity, which has been markedly reduced in the newer high-fidelity variant KpnI-HF (New England Biolabs, #R3142).

Bead trapping and DNA immobilization were carried out as described in the previous section. Once the DNA molecule was tethered between the two beads, the 532 nm excitation laser was set to 10% power.

The workflow for these experiments was as follows. With all valves initially closed and vented, the DNA molecule was positioned at the junction between channel 3 and the protein reservoir. The applied force on the DNA was adjusted to 20 pN, and the pressure was set to 0.8 bar, while keeping all valves closed. A 2D confocal scan was then initiated, followed by opening of channel 6. Immediately thereafter, the relevant reservoir channel (channel 4 or 5) was opened, and the DNA was moved into that reservoir.

For these assays, the reservoir contained 20 µL of KpnI (10 U/µL) in a total volume of 500 µL of NEBuffer™ 2.1 (New England Biolabs, #B7202; discontinued in 2021), composed of 10 mM TrisHCl (pH 7.9), 50 mM NaCl, 10 mM MgCl₂, and 100 µg/mL BSA. Under these conditions, KpnI retains approximately 75% of its full enzymatic activity.

### C-Trap data processing

All data processing was performed using custom-written Python scripts, available at https://github.com/Moreno-HerreroLab/C-TrapDataProcessing. Data in HDF5 files was processed to extract branch photon count profiles from 1D kymographs and branch-length trajectories from 2D confocal scans.

To generate the photon count profiles from kymographs, we first plotted the photon density across the entire kymograph to identify the branch regions. For each branch, a range of 10 to 20-pixel, (0.25-0.5 µm, given a pixel size of 0.025 µm/pixel,) was defined, centred on the branch. The final region was adjusted to match the Gaussian distribution of the photon density for each branch, ensuring that no signal was excluded. Then, photon counts within each area were summed along the DNA-Y-axis, yielding the total branch fluorescence signal for a given time (55 ms/time-line in **Fig. 4** and **Fig. S2**; and 25 ms/time-line in **Fig. S5**). To reduce noise and facilitate interpretation, a Savitzky-Golay filter was applied to the photon count profiles; typically using a 10-point window (or 50-point for long-duration kymographs). Both the filtered and raw photon counts signal were represented for comparison. For clarity, only a subset of the raw data was displayed in plots derived from long kymographs. In such cases, this is noted in the corresponding figure caption.

To extract the apparent branch lengths from the confocal scans, first, the images were averaged across the entire time course to generate a composite image. From the composite image, we defined five ROIs of 25 pixels-width, one baseline containing only the axial DNA, and one for each branch. Then, these ROIs were applied to each frame individually to extract each branch photon counts over time. Photons were summed along the x-axis (DNA axis) to compensate for the lateral movement. Photon profiles were then aligned to their maximum intensity, corresponding to the junction between the branch and the backbone, to cancel the curvature effect of the axial DNA due to the perpendicular flow. Apparent branch lengths were then determined from the profiles using a fluorescence threshold of 20-35 photons, depending on the signal-to-noise ratio. The branch-backbone junction was defined as the zero point, and the axial DNA baseline established the detection limit. Signals below 15 pixels (750 nm) are considered indistinguishable from background and reported as constant length. Branch-length trajectories for the different assays were plotted over time to monitor protein activity.

## Data Availability

Raw and processed data supporting this work are available on Zenodo at ^55^

## Code Availability

Custom Python code used to process raw data and generate the plots presented in this work is available on Github.at https://github.com/Moreno-HerreroLab/C-TrapDataProcessing

## Acknowledgements

We thank Neville Gilhooly and Gemma Fisher from the Dillingham laboratory for their generous gifts of *Bacillus subtilis* AddAB and ParB, respectively.

Research in the F. M-H. and O.L. laboratories was funded through the R&D activities program with reference TEC-2024/TEC-158 and acronym TecNanoBio-CM, granted by the Autonomous Region of Madrid through the “Dirección General de Investigación e Innovación Tecnológica” and by “la Caixa” Foundation under the project code LCF/PR/HR24/52440009- “LncRNAs modulating DNA damage and repair: towards novel therapies for hepatocellular carcinoma”. Research in the FMH laboratory was also supported by grant PID2023-146255NB-I00 from Agencia Estatal de Investigación (AEI/10.13039/501100011033), Ministerio de Ciencia, Innovación y Universidades and co-funded by the European Regional Development Fund (ERDFEU). Research in the O.L. laboratory was also funded by grant PID2023-146110NB-I00 from Agencia Estatal de Investigación (AEI/10.13039/501100011033), Ministerio de Ciencia, Innovación y Universidades and co-funded by the European Regional Development Fund (ERDFEU). O.L. laboratory also had the support from the National Institute of Health Carlos III to CNIO. Research in the MD laboratory was supported by Medical Research Council Grant MR/Y012070/1 and Biotechnology and Biological Sciences Research Council Grant BB/Y004426/1.

## Author contributions

Conceptualization: F.M.-H., S.d.B and C.A.-R.; Optical Tweezers data acquisition, code development and data processing: S.d.B.; DNA preparation: C.A.-R.; Protein materials: A.R.-C., R. A.-B., O.L, C.A.-R. and M.S.D; Writing - original draft: F.M.-H. and S.d.B; Writing - review and editing: F.M.-H, S.d.B and C.A.-R. with input from all authors.

## Supplementary Figures and Tables

**Supp. Fig. S1.**
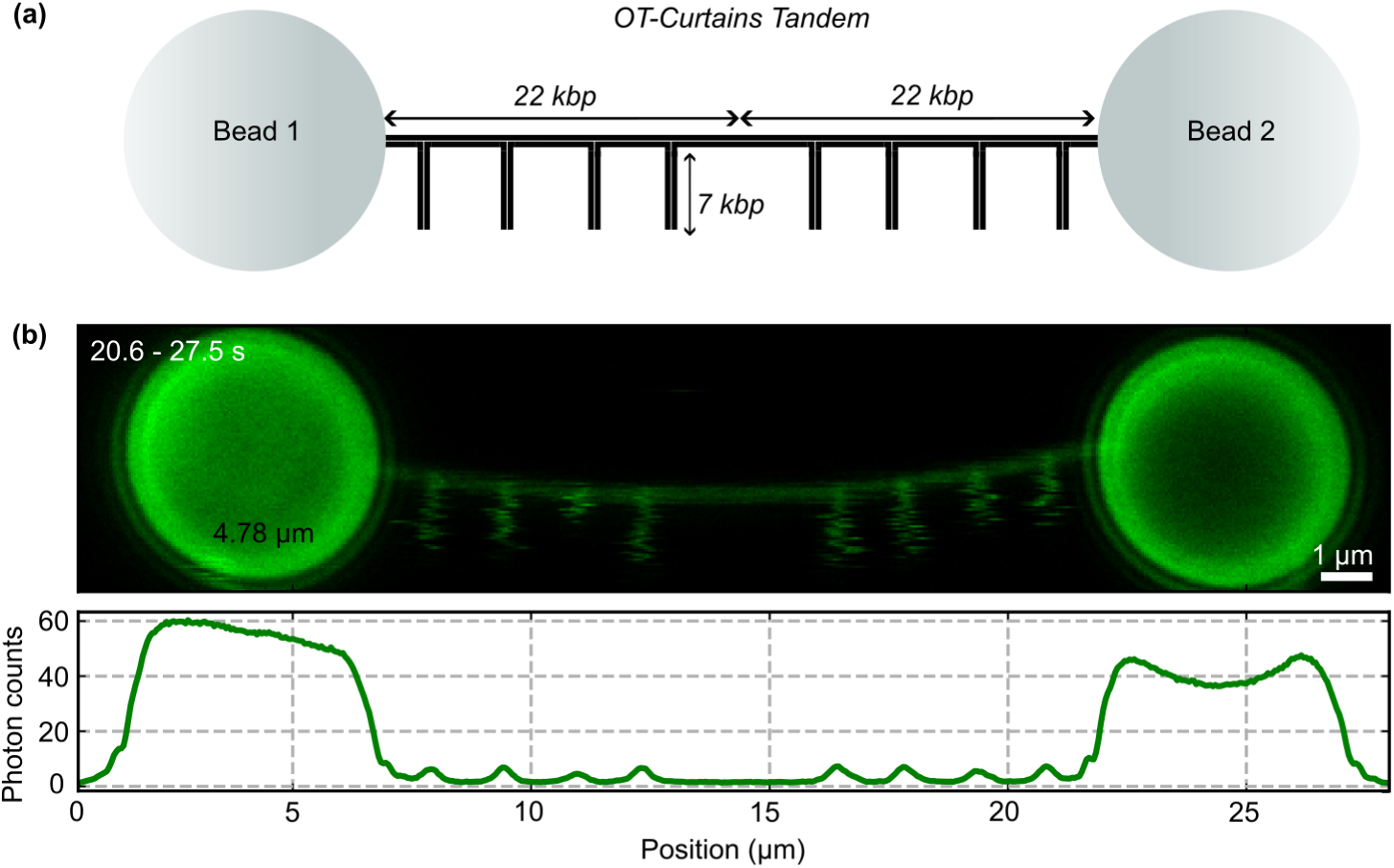
Tandem OT- Curtains. **(a)** Schematics and **(b)** experimental data from a tandem OT-Curtains containing long 7 kbp branches terminating in blunt ends. DNA is labelled with 100 nM Sytox Orange (in green).

**Supp. Fig. S2.**
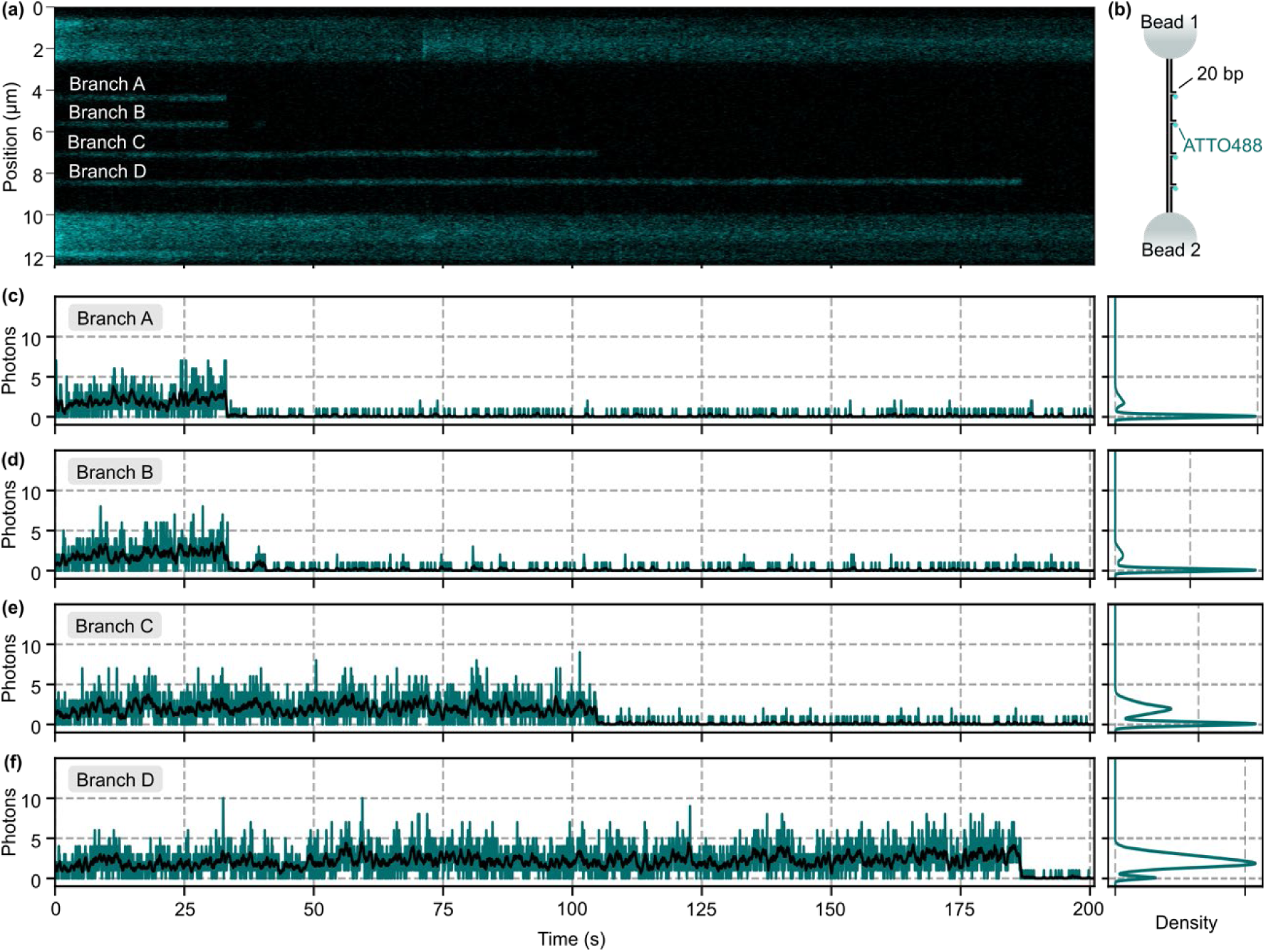
OT-Curtains with short ATTO488-labeled DNA ends. **(a-b)** Kymograph and schematics of ATTO488 labeled OT-Curtains. **(c-f)** Photon counts obtained for areas of 10 pixels around each branch (pixel size of 0.025 µm), for branches A to D, showing photobleaching events. Data corresponds to a single ATTO488 fluorophore. Black lines represent filtered signal (see methods).

**Supp. Fig. S3.**
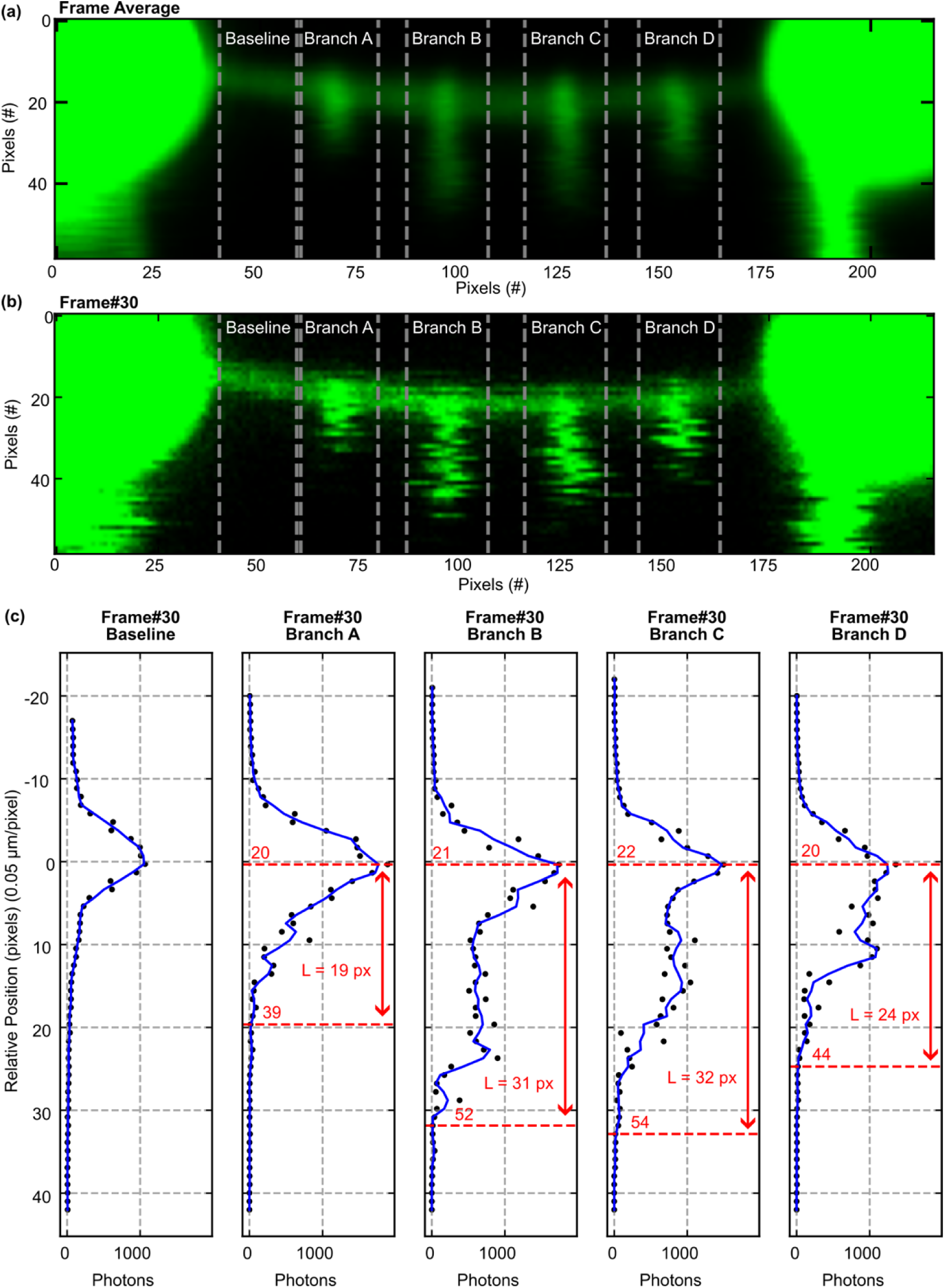
Branch length quantification. **(a)** Average of all frames and defined ROIs for branches and baseline. **(b)** Example of a time frame with ROIs defined in (a). **(c)** Photon counts sum along x-axis.

**Supp. Fig. S4.**
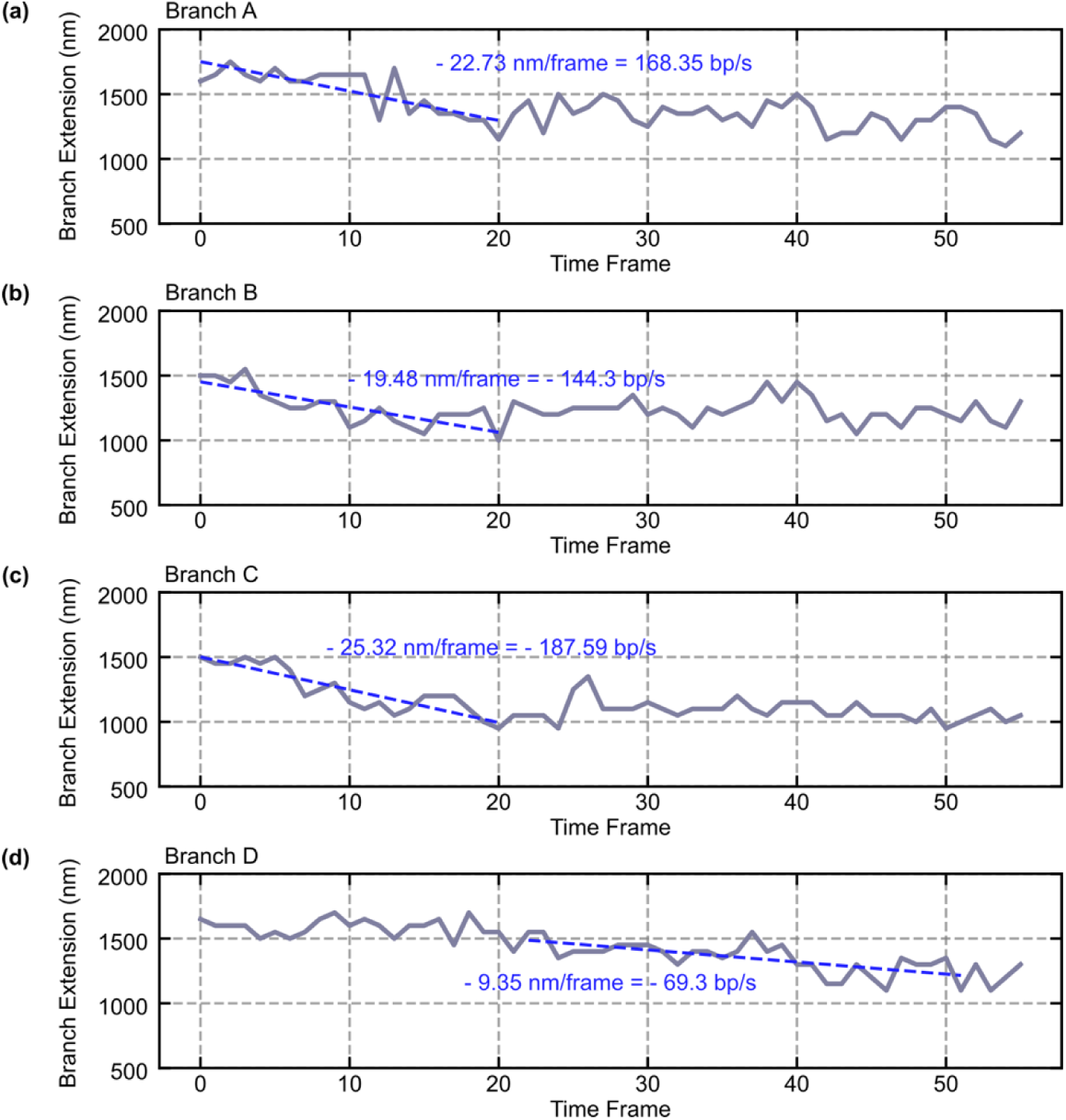
AddAB unwinding/resection rate. **(a-d)** Branch-length trajectories of branches A to D, showing rates for a burst event by AddAB.

**Supp. Fig. S5.**
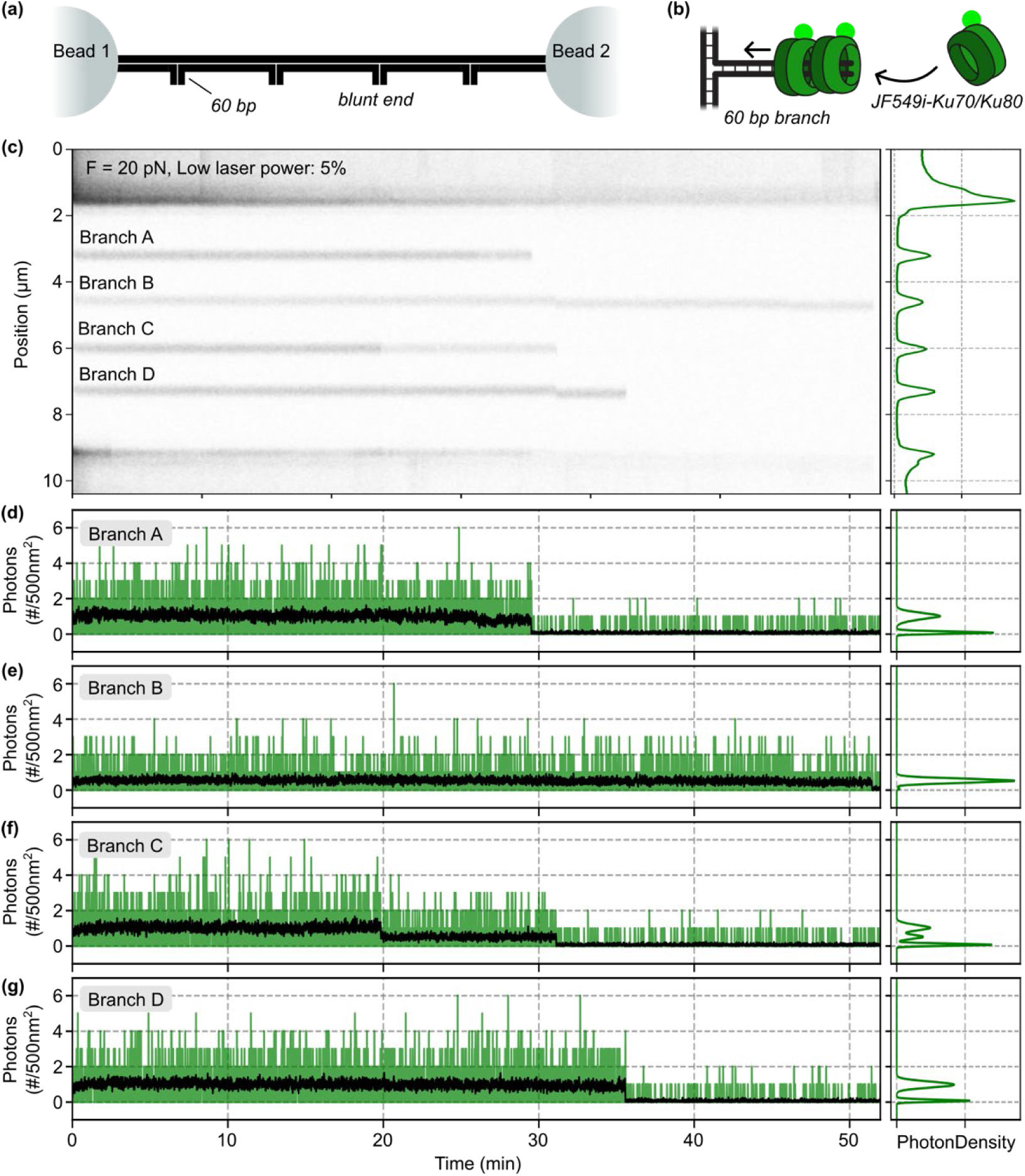
JF549i-Ku remains bound to DNA ends for tens of minutes. **(a-b)** Schematics of OT-Curtains and branches, respectively, used in this assay. **(c)** Kymograph of the OT-Curtains with short 60 bp DNA-ends (unlabeled) in the presence of JF549-labeled Ku (in green). **(d-g)** Photon count profiles from branches A to D.

**Supplementary Table S1.** Reagents.

| Reagent or resource | Source | Identifier |
| --- | --- | --- |
| Bio-16-dUTP | Roche | 11 093 070 910 |
| Bovine Serum Albumin (BSA) | Sigma Aldrich | A7030-100G |
| BssHII Restriction Enzyme | New England Biolabs | R0199S |
| BstBI Restriction Enzyme | New England Biolabs | R0519S |
| DH5α Competent Cells | Thermo Fisher Scientific | 18265-017 |
| DMSO | Sigma Aldrich | 276855-100ML |
| DTT | Sigma Aldrich | 646563-10X.5ML |
| Hellmanex™ III | Merk | Z805939-1EA |
| JF549i-HaloTag ligand | Janelia Fluor® Dyes | --- |
| KpnI Restriction Enzyme (non-HF) ** | New England Biolabs | R0142 |
| NEBuffer™ 2.1 ** | New England Biolabs | B7202 |
| Nt.BbvCI Nicking Enzyme | New England Biolabs | R0632S |
| PciI Restriction Enzyme | New England Biolabs | R0655S |
| Phusion High-Fidelity DNA Polymerase | Thermo Fisher Scientific | F-553S |
| Qdot™ 525 Streptavidin Conjugate | Thermo Fisher Scientific | Q10143MP |
| QIAprep Spin Miniprep Kit | QIAGEN | 27106 |
| QIAquick Gel Extraction Kit | QIAGEN | 28706 |
| QIAquick PCR Purification Kit | QIAGEN | 28106 |
| rSAP Phosphatase | New England Biolabs | M0371S |
| Sall Restriction Enzyme | New England Biolabs | R0138S |
| Streptavidin Coated Polystyrene Particles (1.5-1.9 μm) | Spherotech, Inc. | SVP-15-5 |
| Superdex 200, 10/300 Increase | Cytiva | GE17-5175-01 |
| Synthetic Oligonucleotides, HPLC purified | Sigma Aldrich | --- |
| T4 DNA Ligase | New England Biolabs | M0202S |
| T4 Polynucleotide Kinase | New England Biolabs | M0201S |
| Trolox | Sigma Aldrich | 238813-1G |

(\*\* discontinued product)

**Supplementary Table S2.**
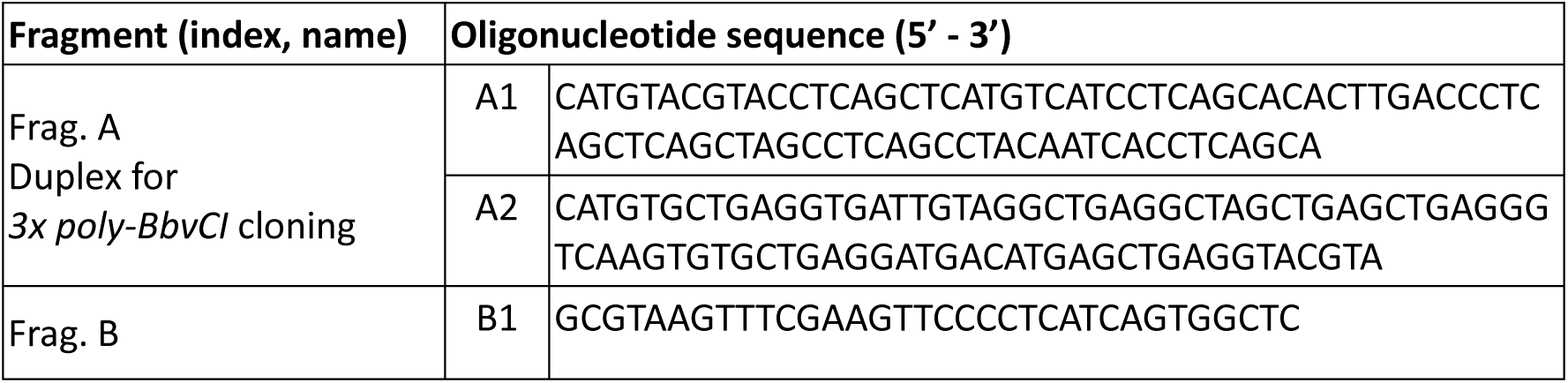
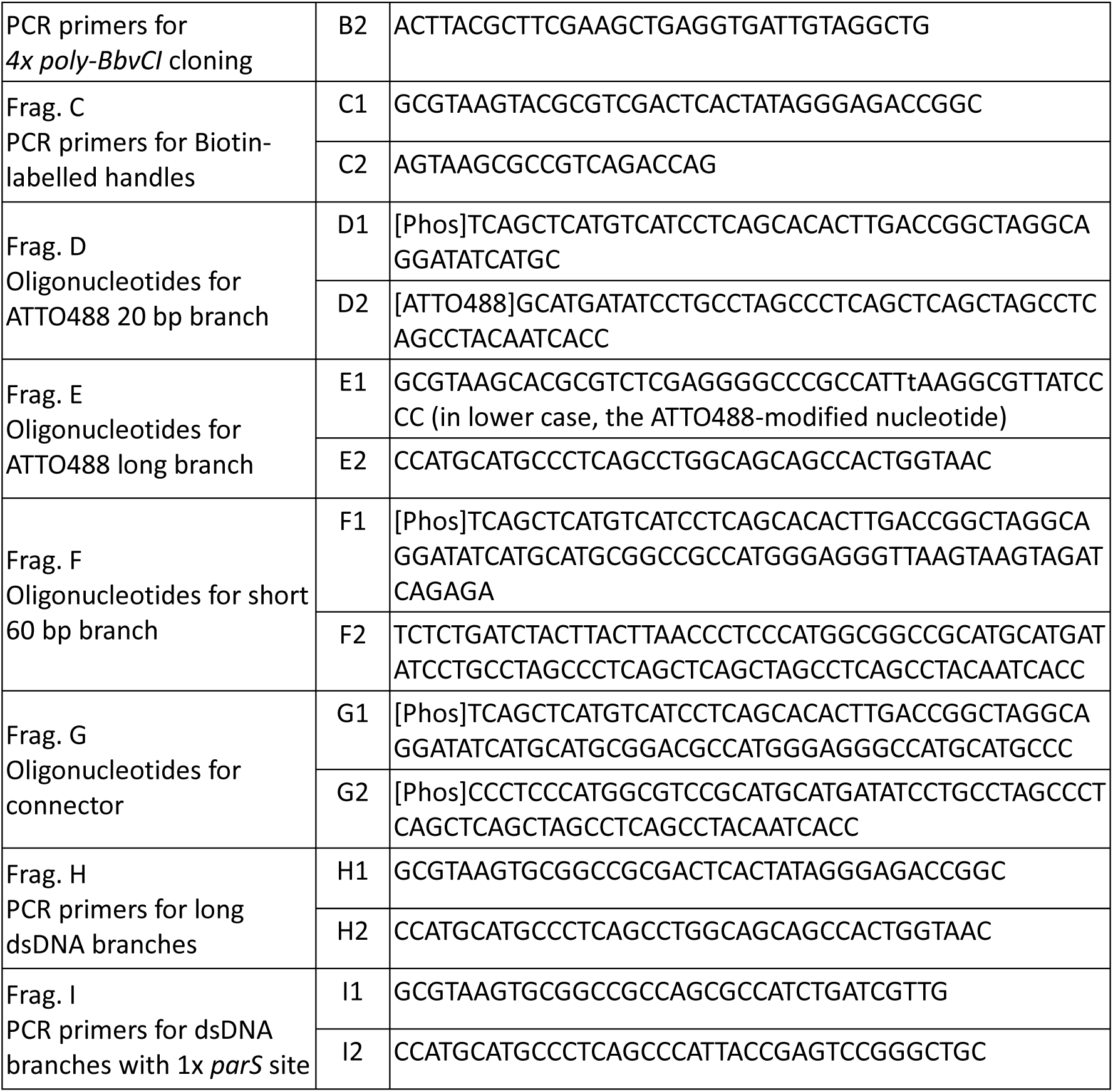
Oligonucleotides.

**Supplementary Table S3.**
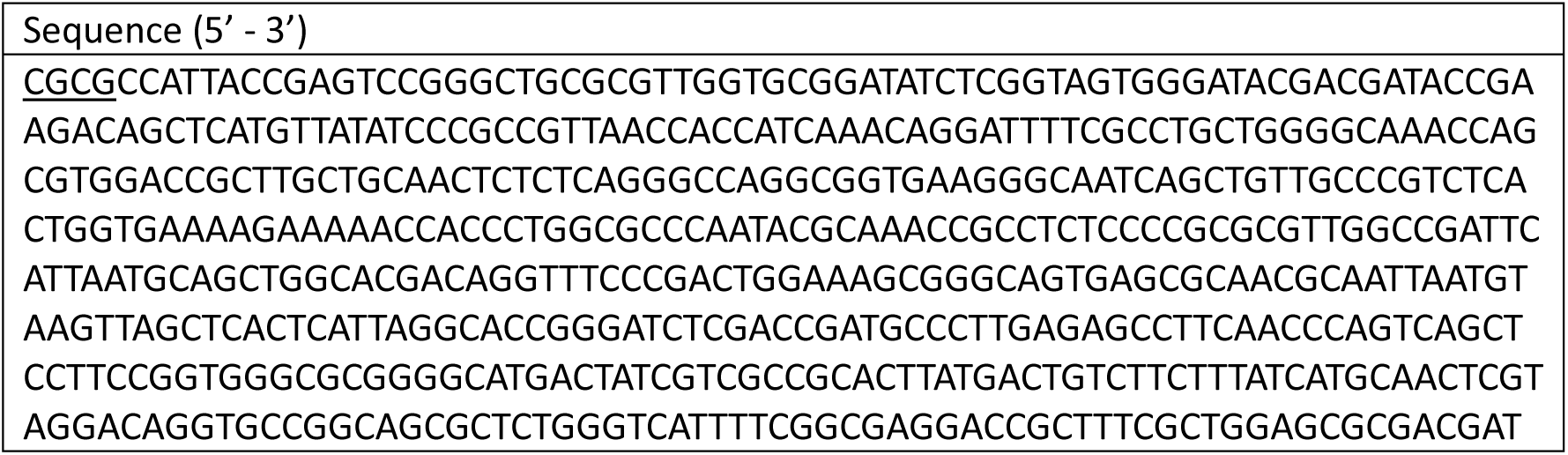
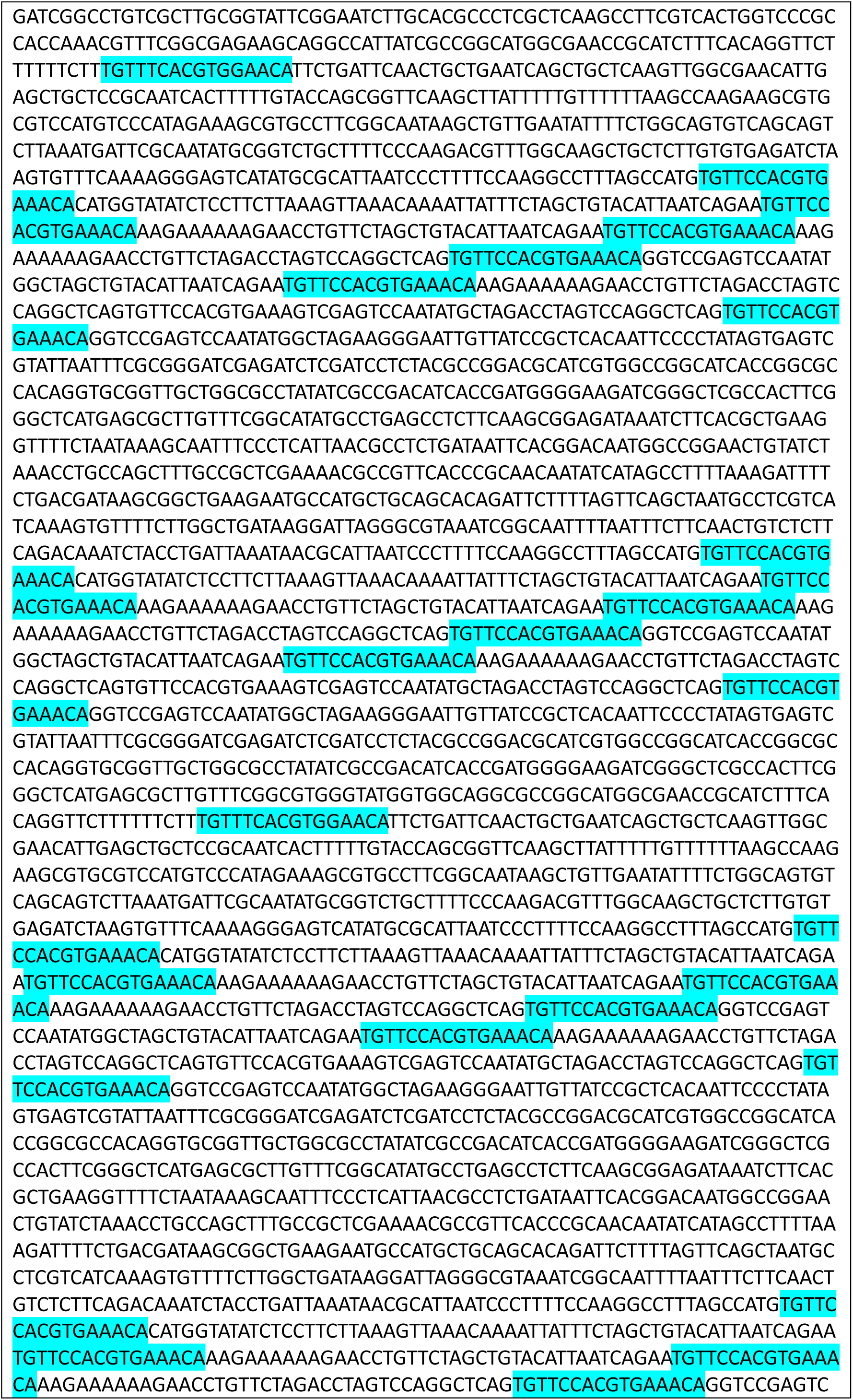
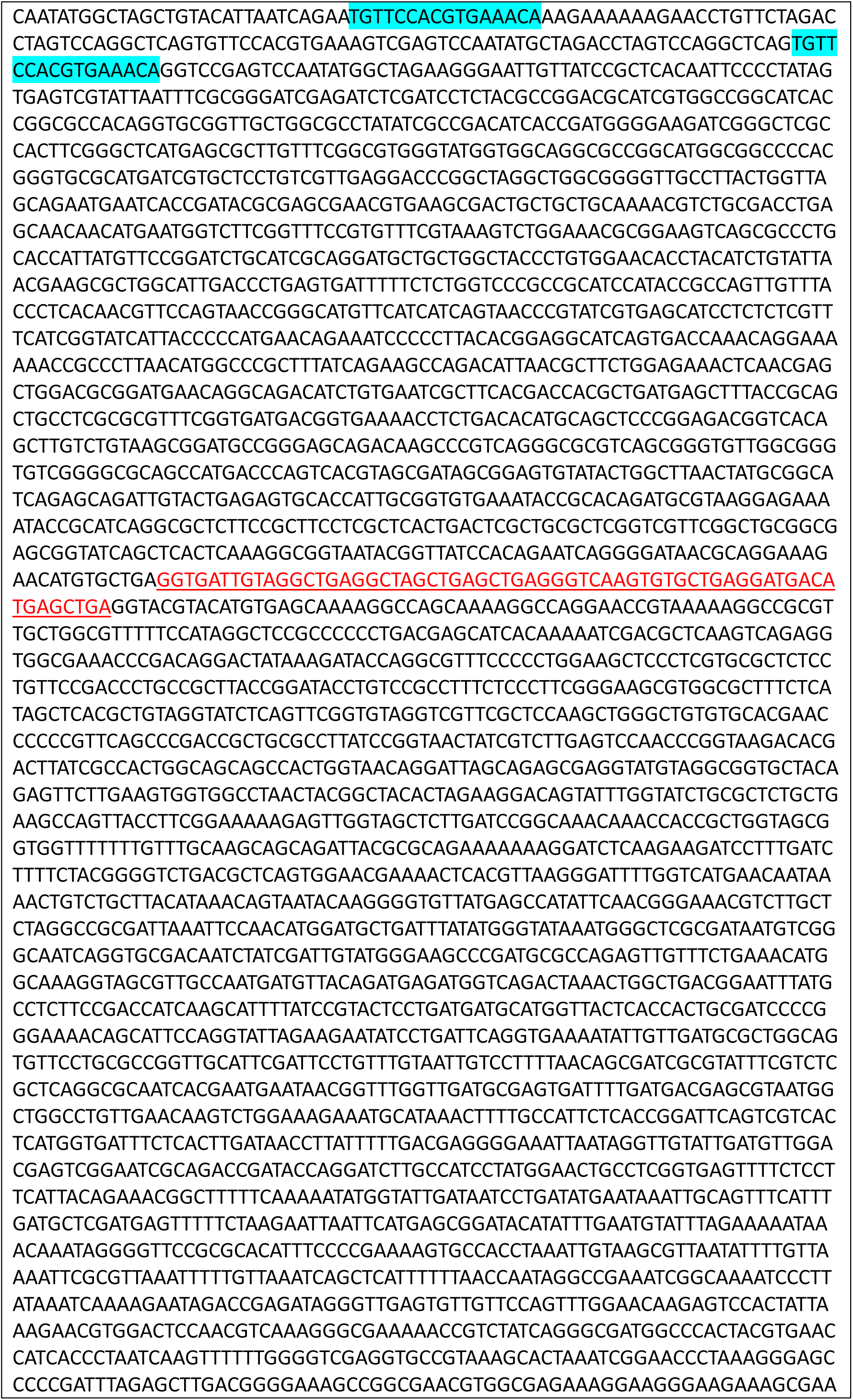
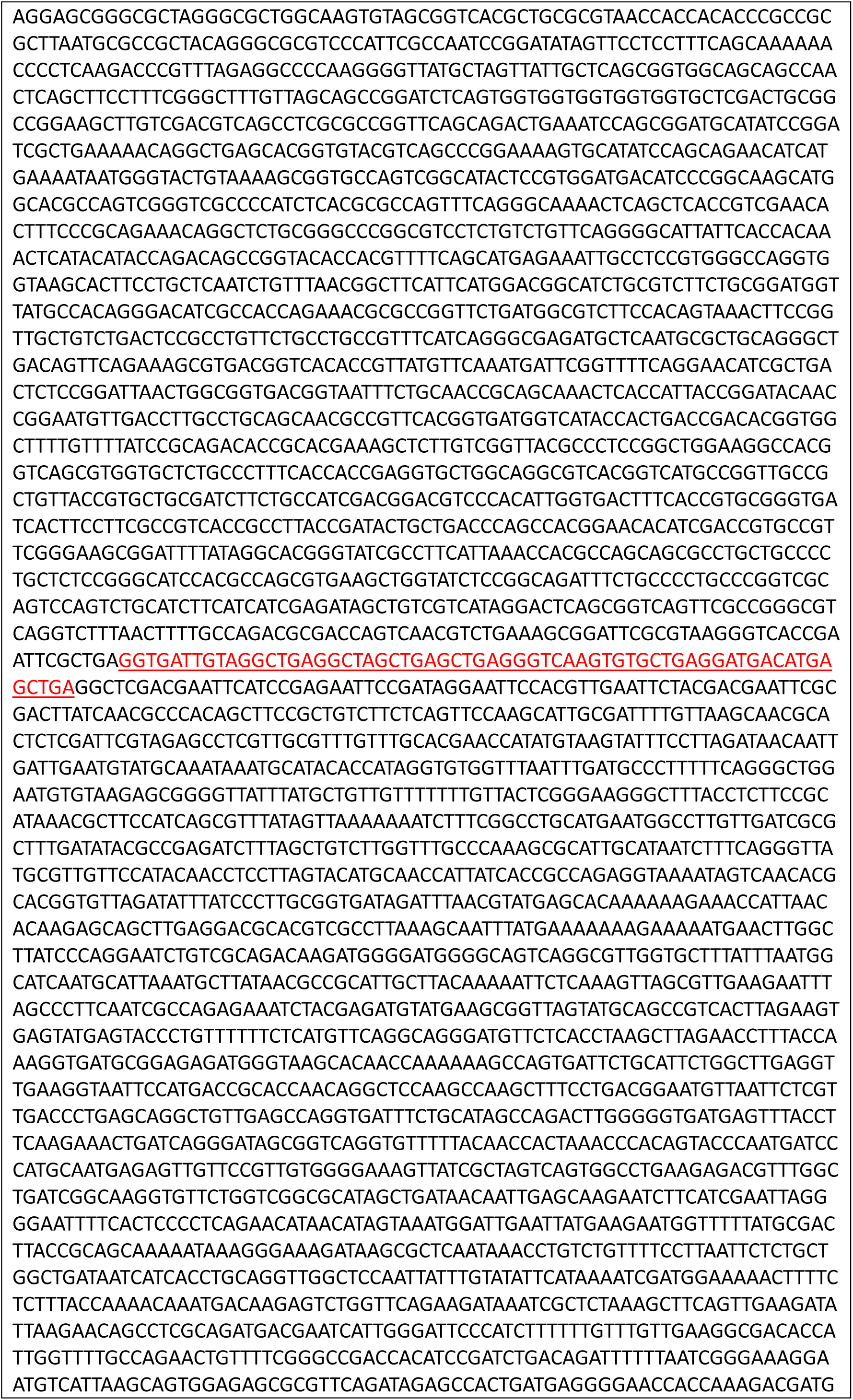
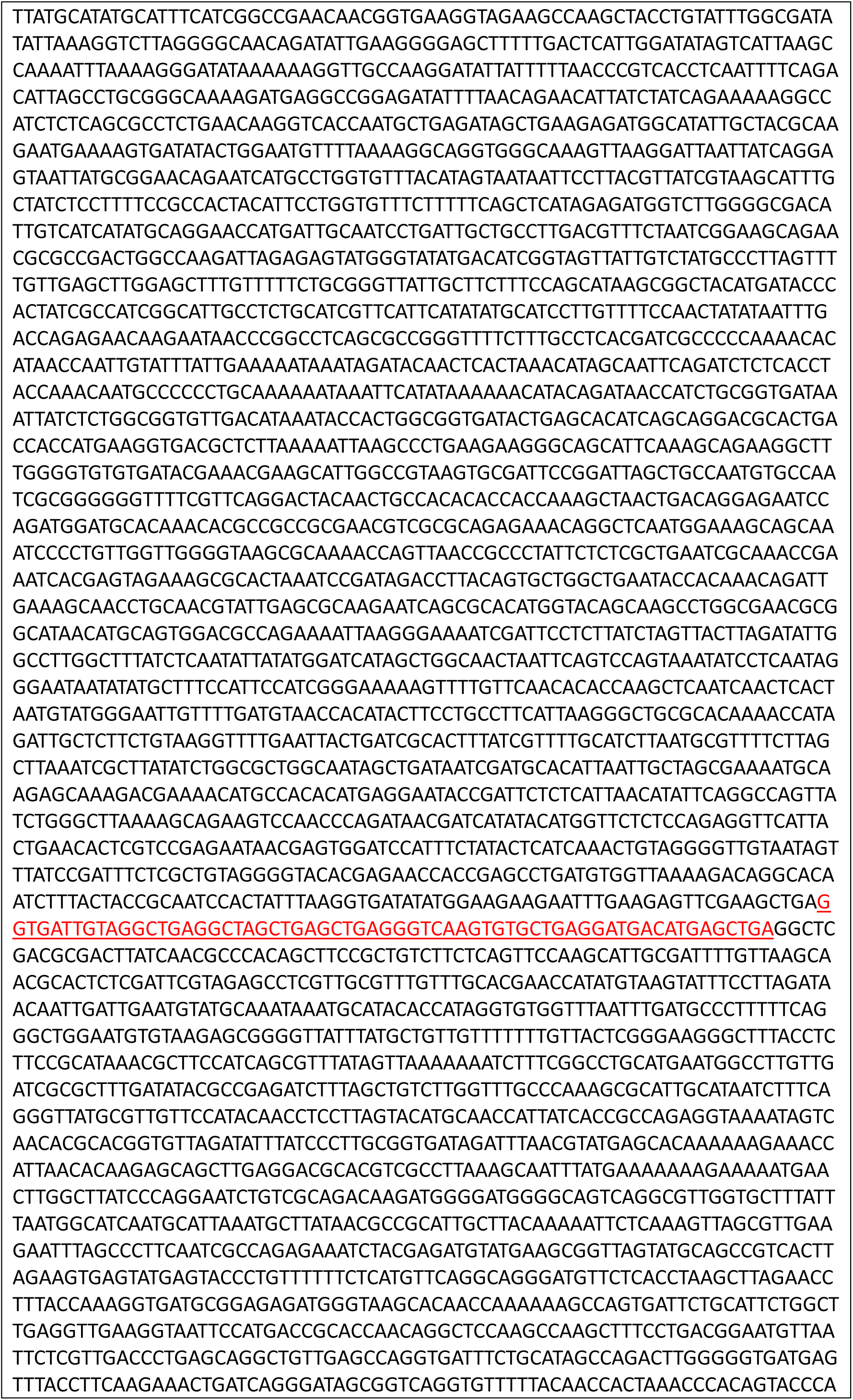
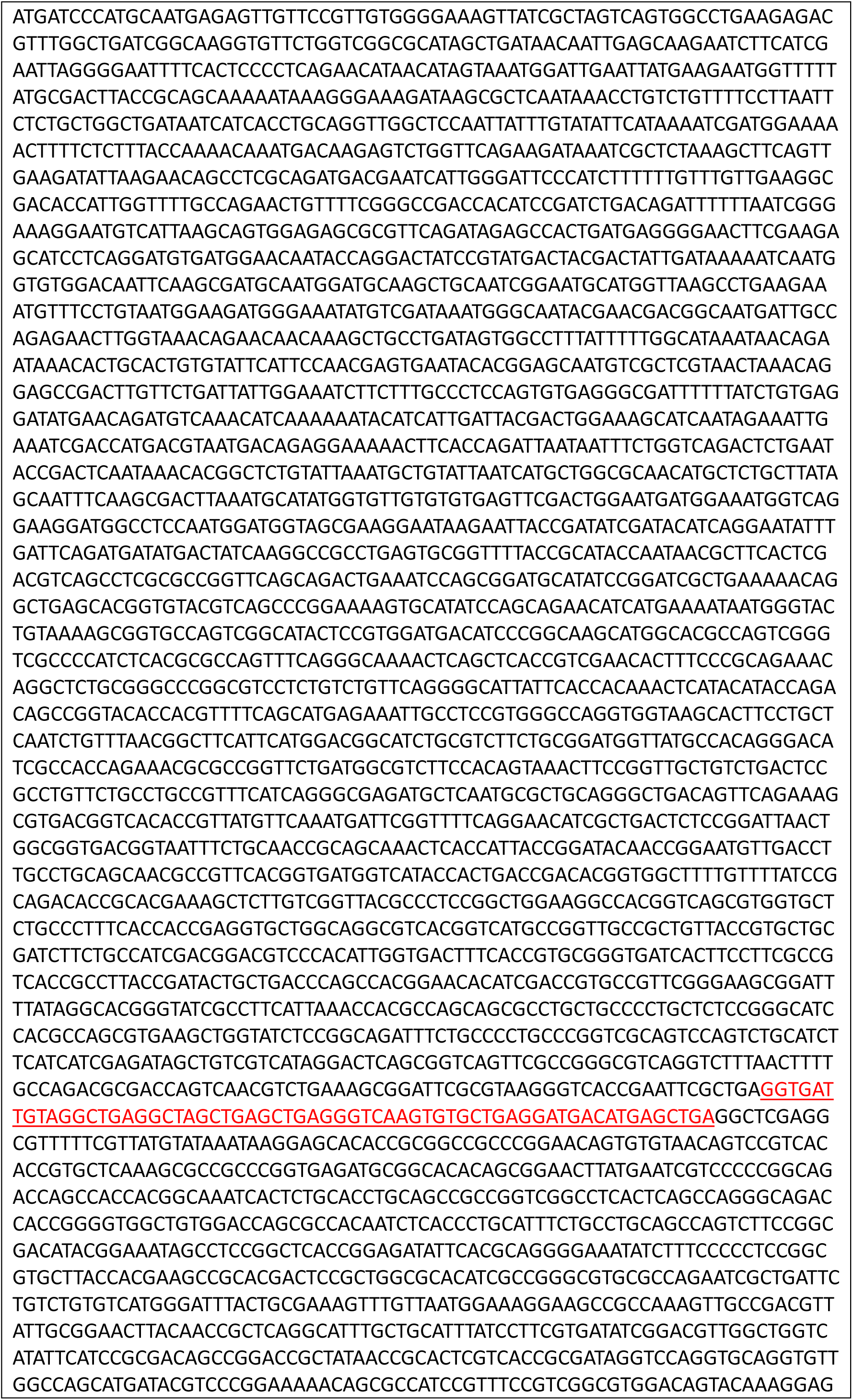
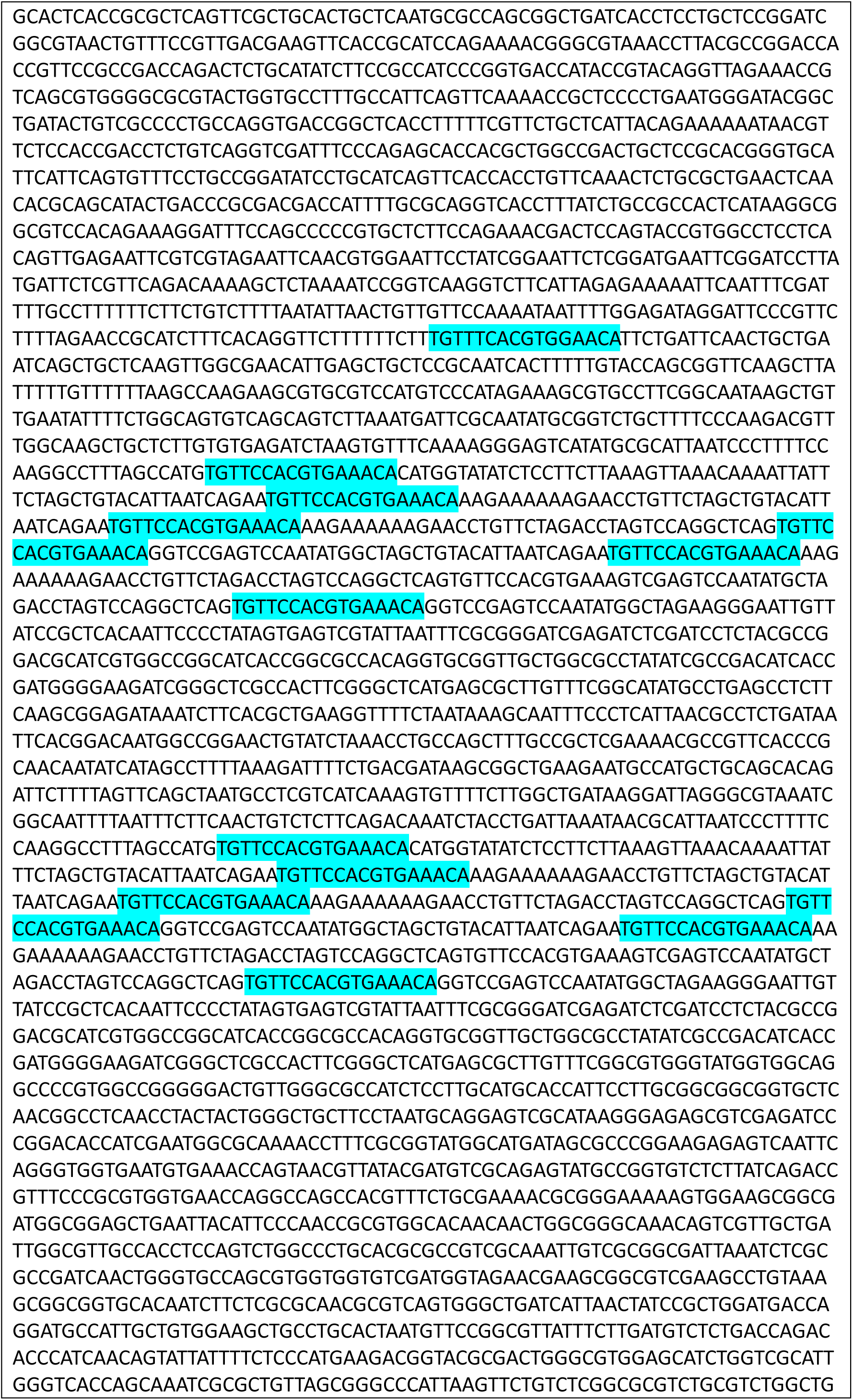
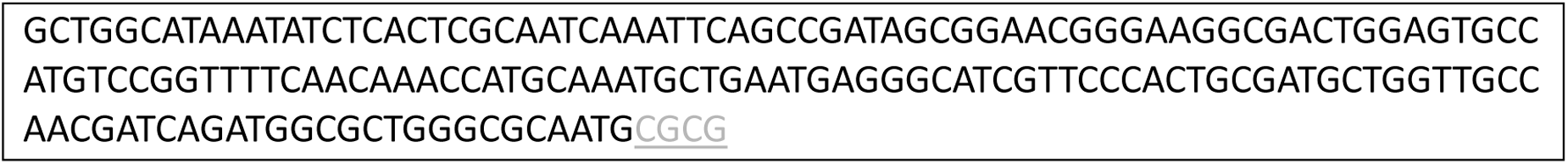
OT-Curtains backbone with 4x ssDNA gaps (22,138 bp). Regions underlined in red indicate the 63 nt single-stranded gaps generated after digestion with the nicking enzyme Nt.BbvCI, followed by strand denaturation. Highlighted in blue are the *parS* sites. Underlined black letters denote 5’ overhangs on one strand, and underlined grey letters denote 5’ overhangs on the complementary strand.

**Supplementary Table S4.**
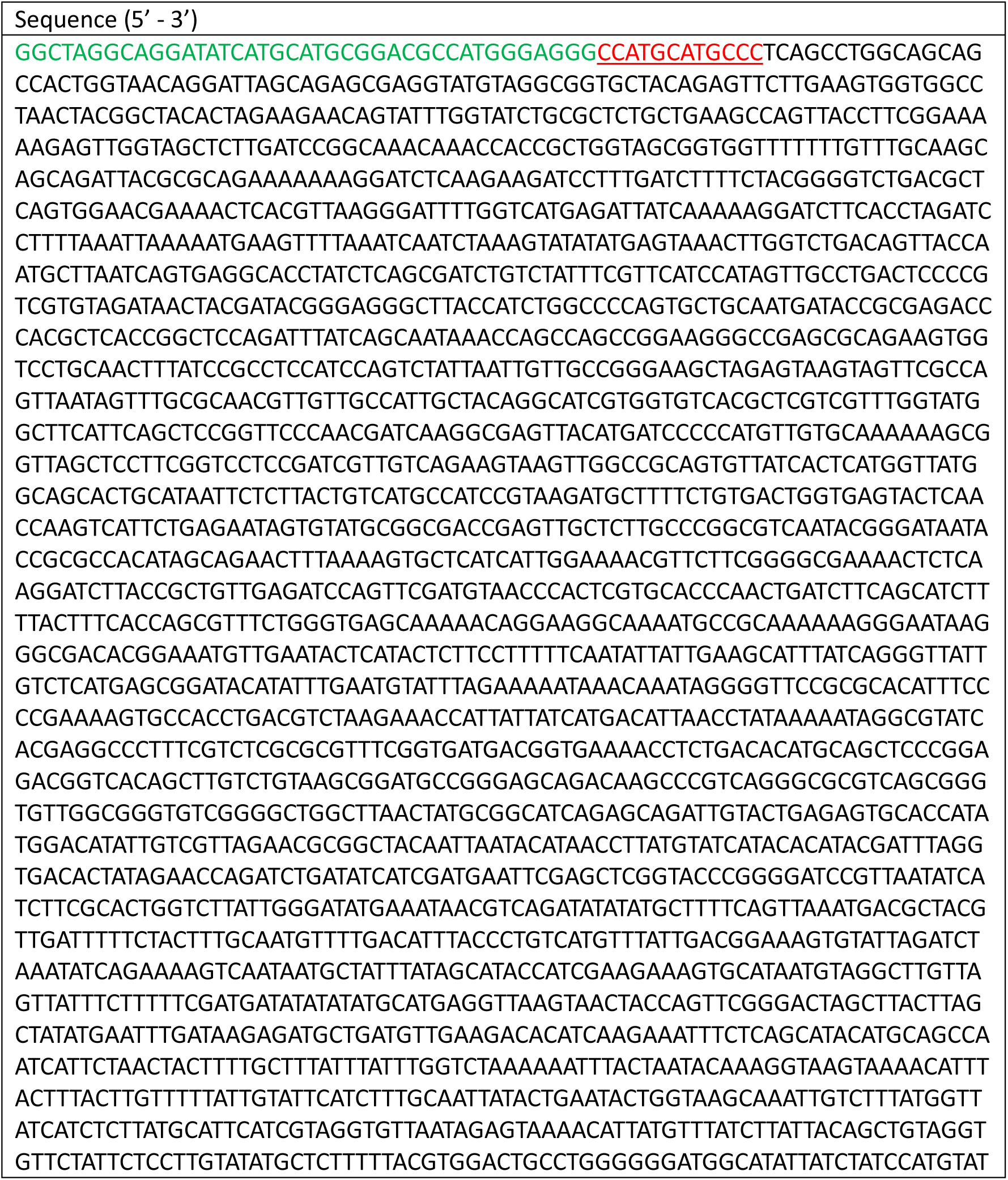
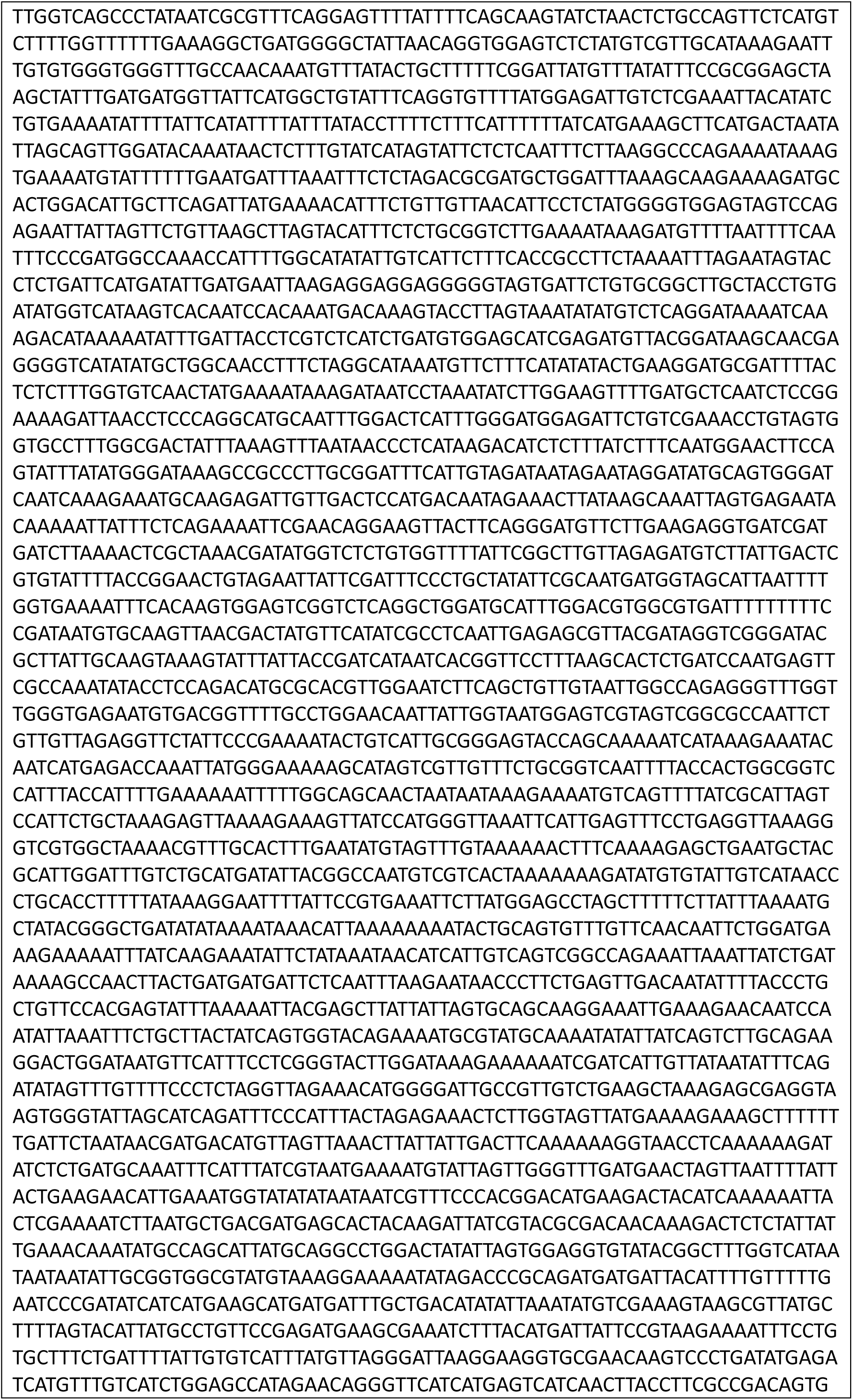
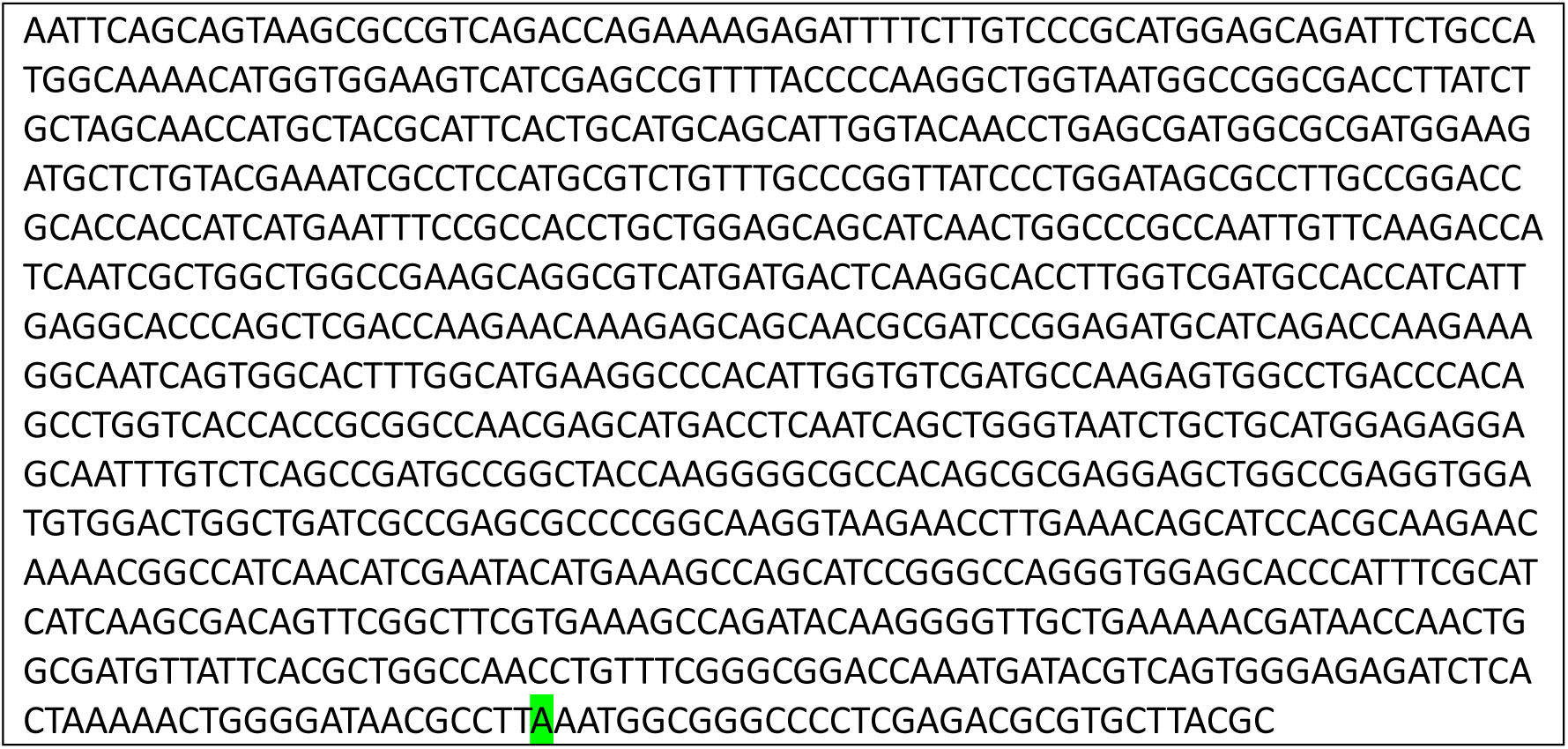
Long dsDNA ATTO488-labelled branches (7,148 bp). Regions in green correspond to the 40 bp stem of the connector. Underlined in red are the 12 bp generated by annealing the long DNA fragment to the connector. The highlighted green position indicates the site of the ATTO488 fluorophore modification located on the complementary strand.

**Supplementary Table S5.**
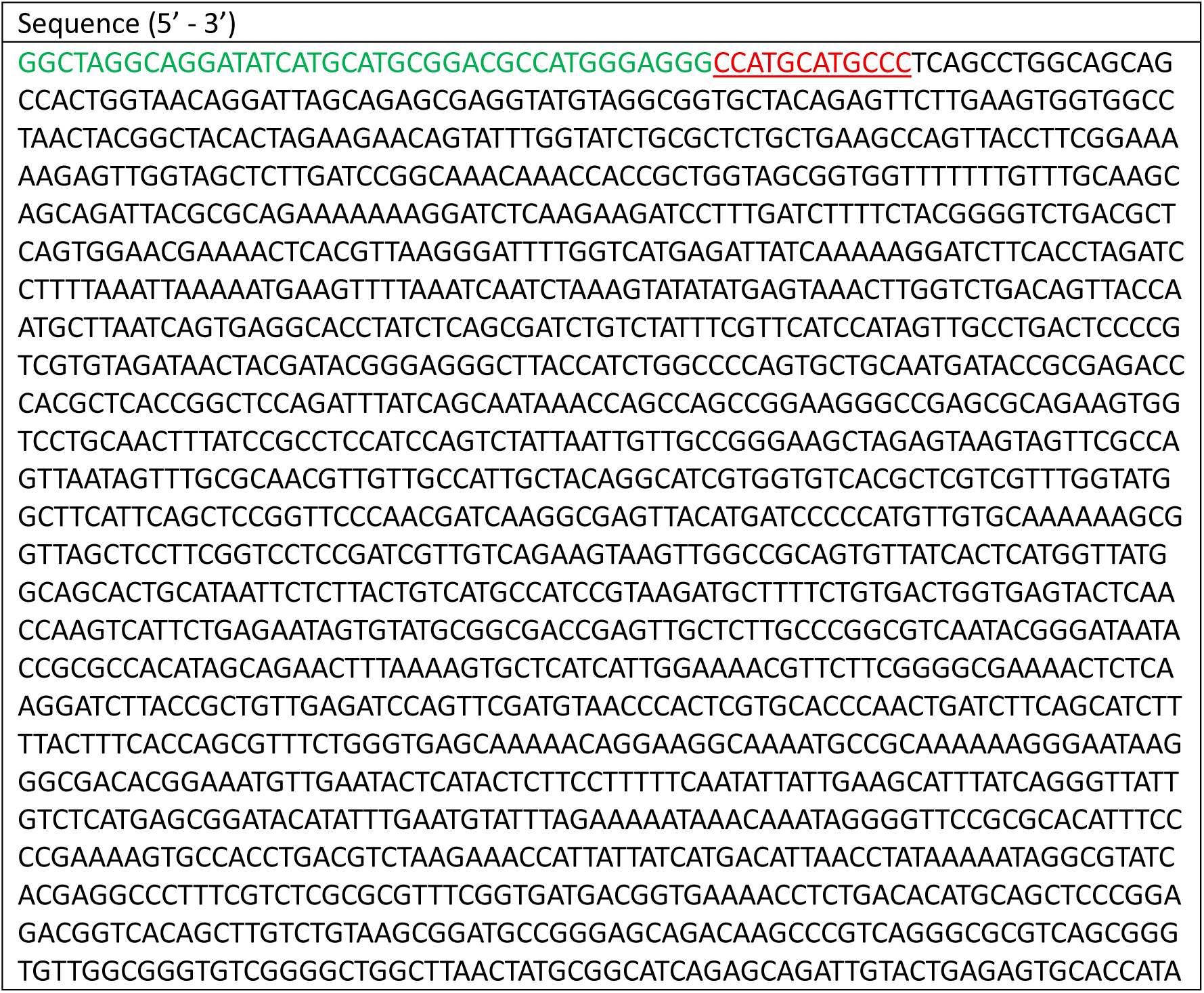
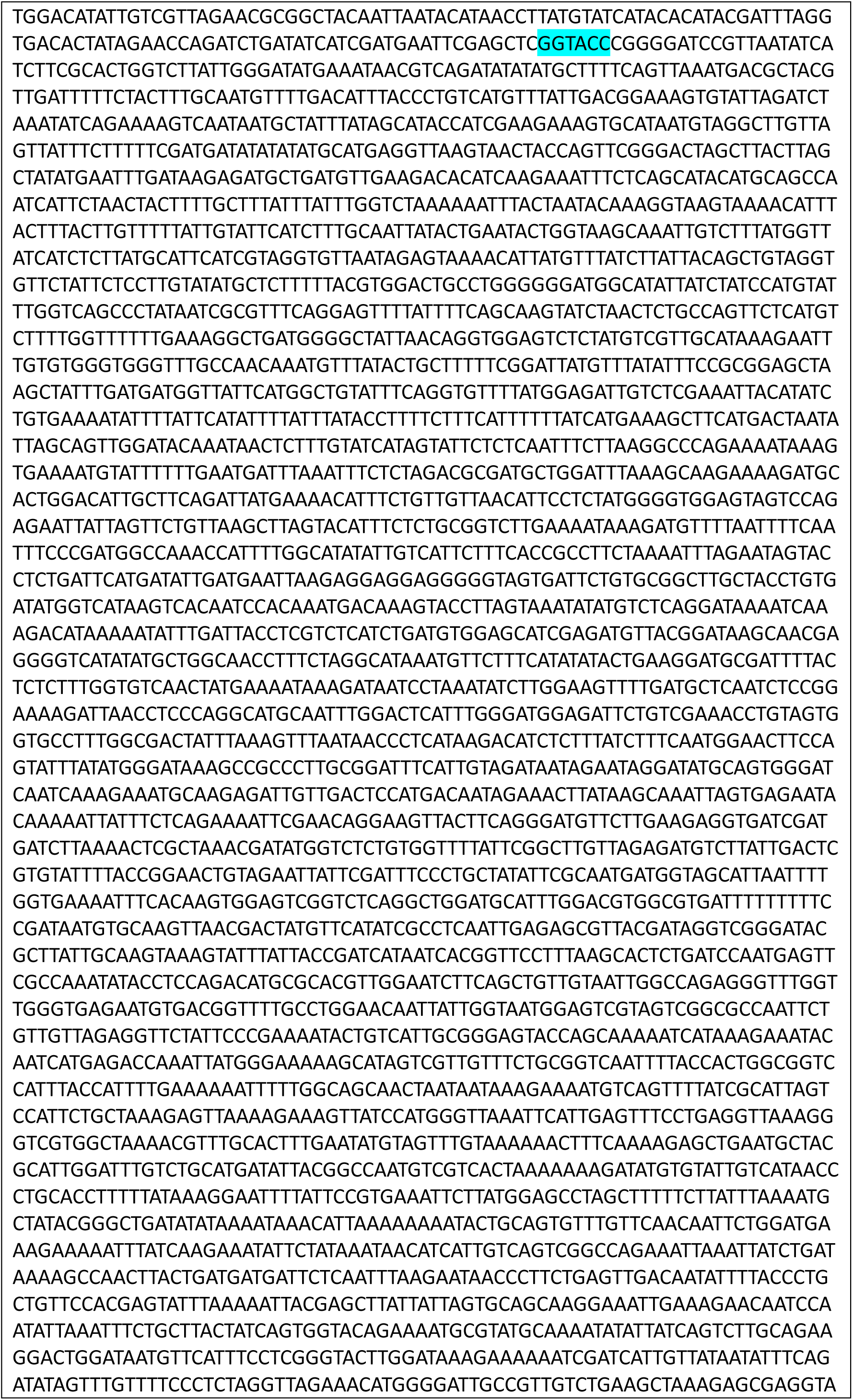
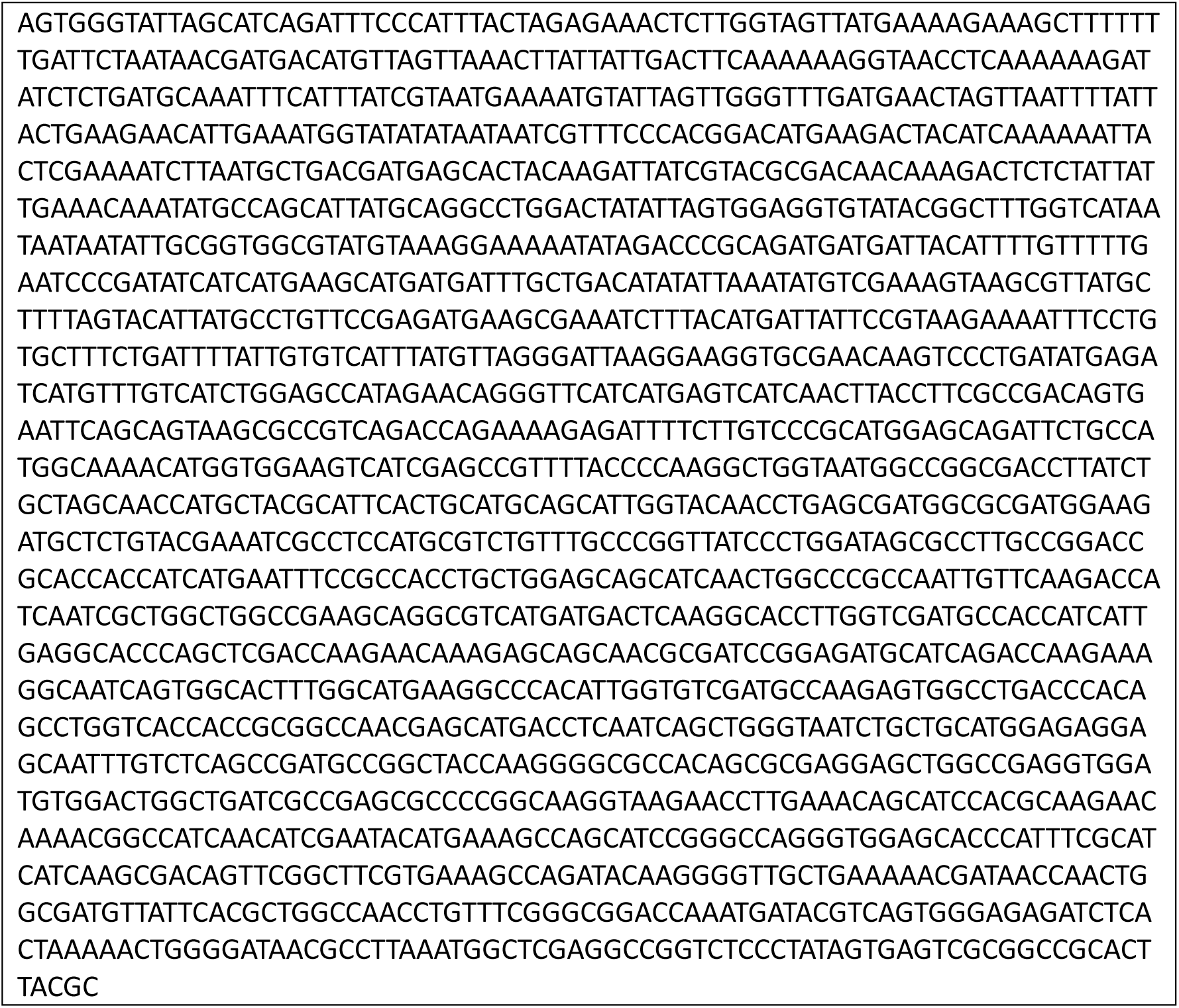
Long dsDNA branches (7,165 bp). Regions in green correspond to the 40 bp stem of the connector. Underlined in red are the 12 bp formed by annealing the long DNA fragment to the connector. The highlighted blue sequence marks the KpnI recognition site.

**Supplementary Table S6.**
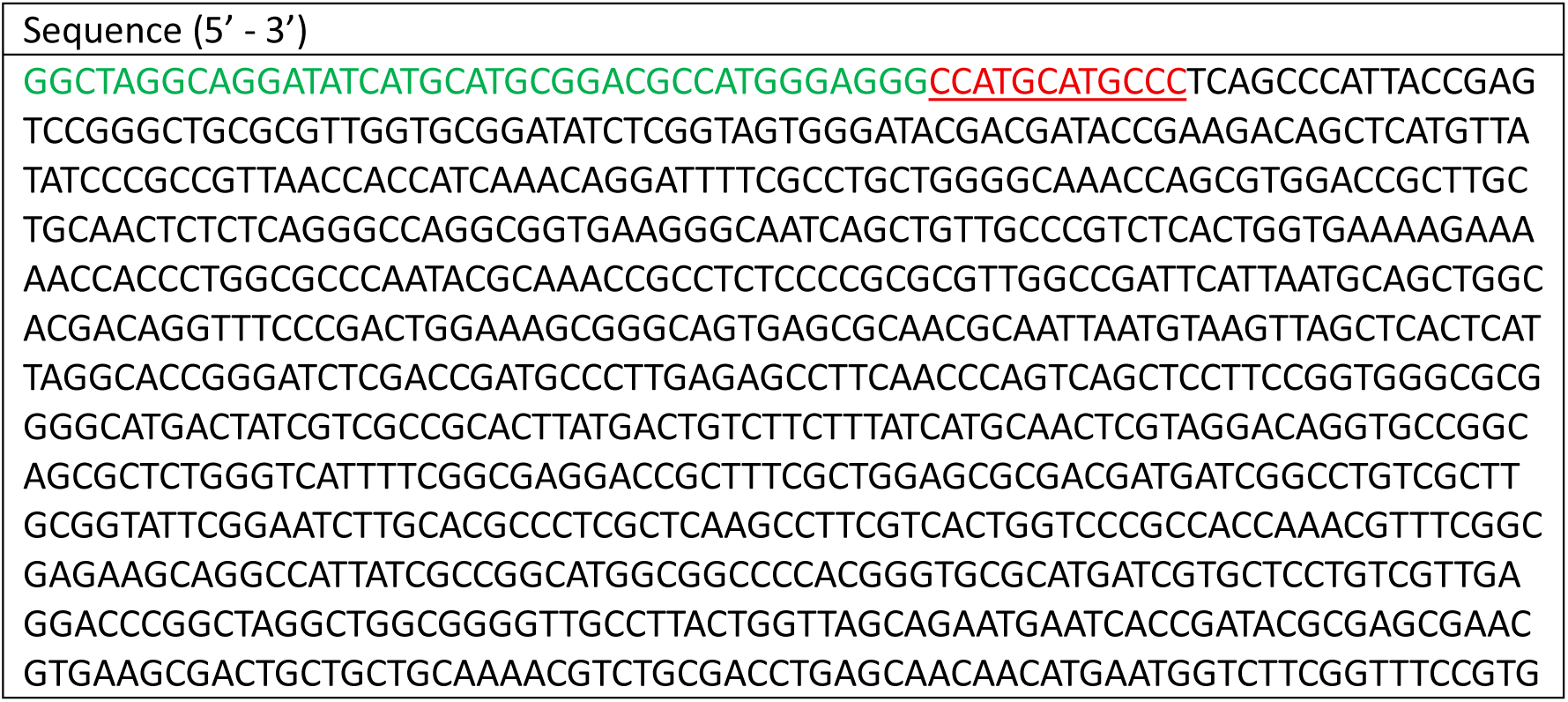
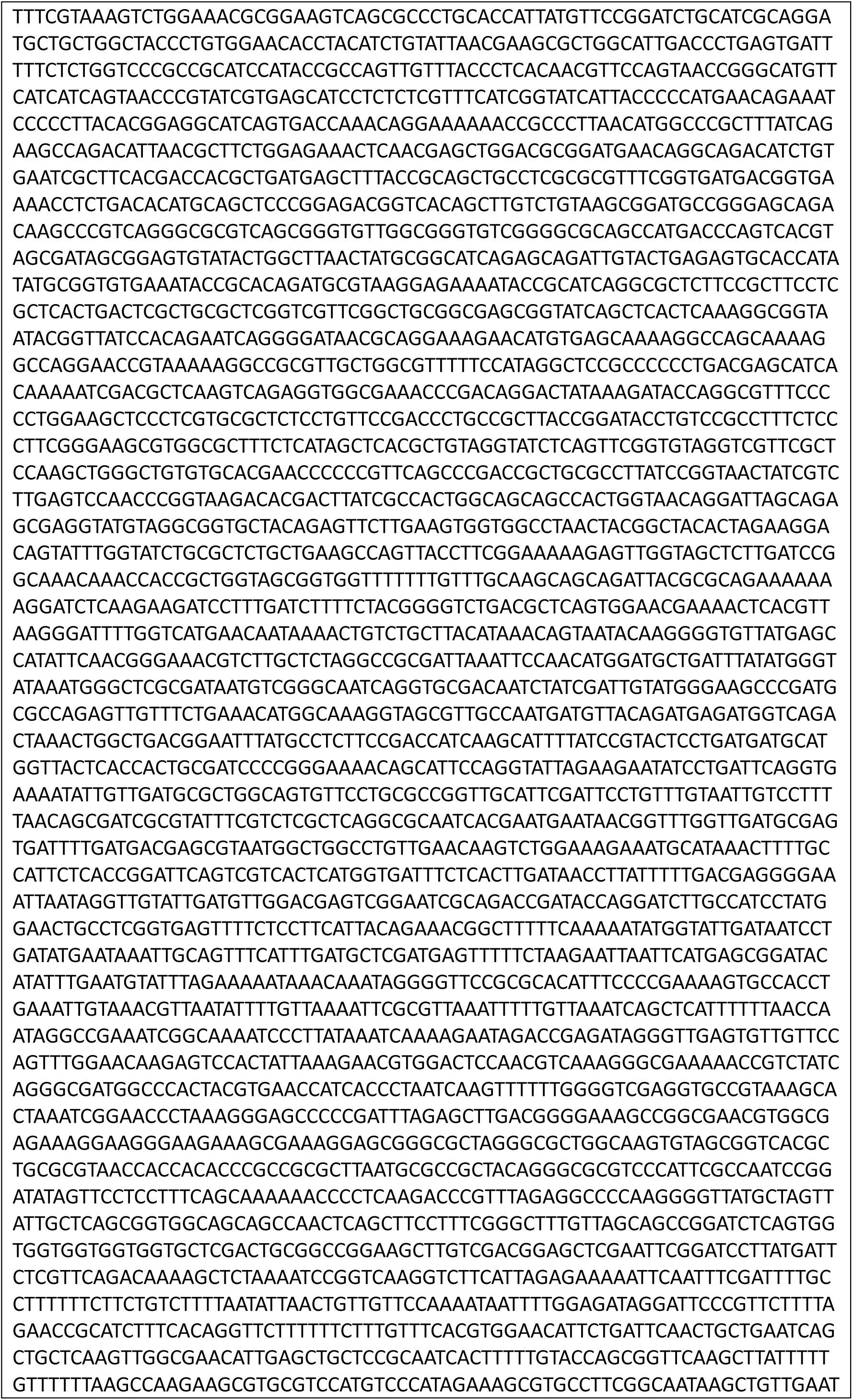
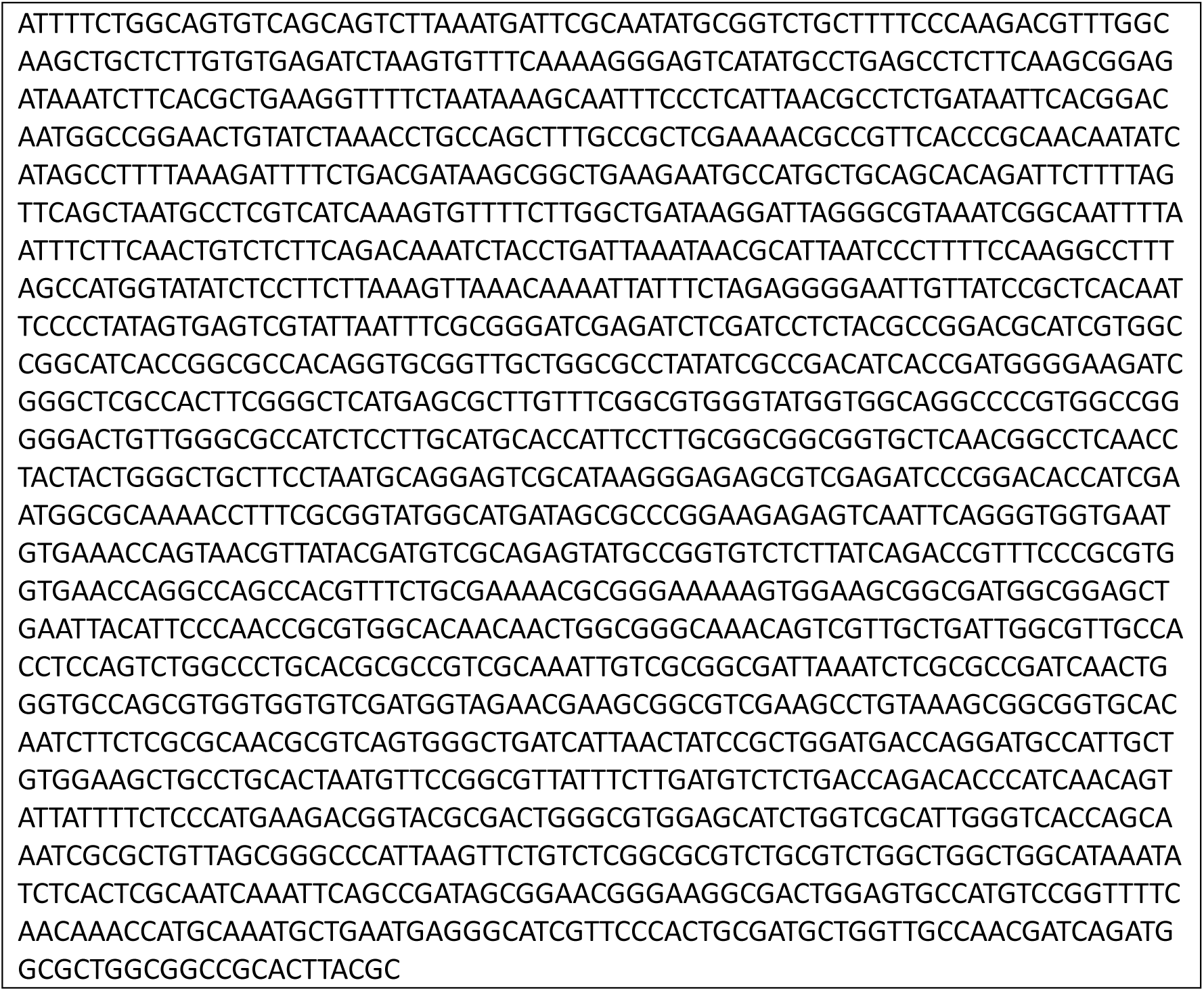
Long dsDNA branches containing one parS site (6,182 bp). Regions in green correspond to the 40 bp stem of the connector. Underlined in red are the 12 bp formed by annealing the long DNA fragment to the connector. The highlighted blue sequence denotes the single *parS* site.

